# Spondweni virus infection in pregnant rhesus macaques causes placental pathology without apparent fetal harm

**DOI:** 10.64898/2026.08.10.743875

**Authors:** Hunter J. Ries, Livia Romanov, Maria C. Charles, Chelsea M. Crooks, Carson DePagter, Alex Richardson, Grace A. VanSleet, Andrea M. Weiler, Jens C. Eickhoff, Katharina S. Stewart, Leandro B. C. Teixeira, Eric Peterson, Michele Schotzko, Heather A. Simmons, Jenna R. Rosinski, Lauren E. Raasch, Anna S. Jaeger, Elaina R. Razo, Emma L. Mohr, David H. O’Connor, Christina M. Newman, Matthew T. Aliota, Thomas C. Friedrich

**Affiliations:** Department of Pathobiological Sciences, University of Wisconsin–Madison, Madison, Wisconsin, USA; Wisconsin National Primate Research Center, University of Wisconsin–Madison, Madison, Wisconsin, USA; Department of Biostatistics and Medical Informatics, University of Wisconsin–Madison, Madison, Wisconsin, USA; Department of Obstetrics and Gynecology, University of Wisconsin–Madison, Madison, Wisconsin, USA; Department of Pathology and Laboratory Medicine, University of Wisconsin–Madison, Madison, Wisconsin, USA; Department of Veterinary and Biomedical Sciences, University of Minnesota, Twin Cities, St. Paul, Minnesota, USA; Department of Pediatrics, University of Wisconsin–Madison, Madison, Wisconsin, USA; Department of Medical Microbiology and Immunology, University of Wisconsin–Madison, Madison, Wisconsin, USA

**Keywords:** Spondweni virus, Zika virus, rhesus macaques, non-human primates, pregnancy, placental pathology, fetal growth

## Abstract

The 2015–2016 Zika virus (ZIKV) epidemic revealed the potential of flaviviruses to emerge rapidly, cause severe disease, and affect pregnancy outcomes. In 2016, Spondweni virus (SPOV), the closest known relative of ZIKV, was detected in mosquitoes in Haiti, suggesting it may also have the potential to emerge in the Western Hemisphere. The risks that close relatives of ZIKV pose to pregnant individuals are not well understood. Previously, we showed that SPOV can cause fetal demise, placental pathology, and vertical transmission in a mouse model. Here we report SPOV’s pathogenic potential in pregnant rhesus macaques. We inoculated four macaques with SPOV at gestational day 30 (early first trimester) and compared their viral loads and fetal outcomes with those of macaques infected in the first trimester with either African-lineage ZIKV (ZIKV-DAK) or an Asian-lineage ZIKV isolate from Puerto Rico (ZIKV-PR) in previous studies. Plasma viremia persisted 10–31 days in SPOV-inoculated dams, whereas viremia resolved within 10–17 days for ZIKV-DAK and 5–52 days for ZIKV-PR. Cesarean deliveries near term (gestational day 152–157) revealed no demise, premature birth, or gross abnormalities in fetuses of dams inoculated with SPOV or ZIKV-PR. In contrast, under near-identical conditions, all ZIKV-DAK-inoculated dams experienced fetal demise between 12 and 20 days post-inoculation. At cesarean section, we did not detect SPOV RNA above the limit of detection in maternal (e.g., spleen, liver), placental, or fetal tissues, in contrast to previous findings with ZIKV-PR. Histological analysis revealed mononuclear/lymphohistiocytic villitis in all placentas of SPOV-exposed macaques, along with other pathological changes in individual placentas. Our findings suggest that SPOV infection of macaques in early pregnancy may result in placental pathology without overt fetal harm. Our results suggest that flaviviruses in the Spondweni serocomplex, which includes ZIKV and SPOV, may vary in their pathogenic potential during pregnancy.

**Author Summary:** Zika virus (ZIKV) can cause fetal harm. Does this risk extend to its closest known relative, Spondweni virus (SPOV)? Should SPOV circulate in humans, what risks would it pose in pregnancy? SPOV can injure fetuses in immunocompromised mice, but the physiology of pregnancy in mice differs greatly from that of humans. We therefore infected pregnant rhesus macaques with SPOV during early gestation and compared maternal viremia, placental pathology, and fetal outcomes with macaques infected with African- or Asian-lineage ZIKVs at the same gestational age. All fetuses survived to near-term pregnancy, fetal tissues were negative for SPOV RNA, and fetal growth tracked within expected ranges. Nonetheless, all SPOV-exposed pregnancies showed placental injury, including mononuclear/lymphohistiocytic villitis and maternal vascular malperfusion. Despite the absence of detectable SPOV RNA in fetal tissues, SPOV RNA persisted at term in maternal-fetal interface tissues in two of four animals. These data indicate placental injury without detectable vertical transmission in this translational model. Our results suggest that SPOV and ZIKV display a wide range of risks to the developing fetus. Identifying viral and host factors that increase the potential for fetal harm will be important for assessing risks posed by emerging viruses in this family.

## Introduction

Zika virus (ZIKV), dengue virus (DENV), and yellow fever virus (YFV) are closely related orthoflaviviruses that cause hundreds of millions of infections annually and place half of the world’s population at risk of infection [1–4]. As climate change expands the geographic range of their primary mosquito vectors, *Aedes aegypti* and *Aedes albopictus*, these flaviviruses pose a growing global health threat [1,5]. Despite their genetic similarity, these viruses exhibit markedly different pathogenic profiles: ∼5% of symptomatic DENV infections progress to severe dengue that can be life-threatening [6–8], ∼12% of YFV infections enter a severe “toxic” phase in which approximately half of cases are fatal [9,10], and ZIKV infection during the first trimester of pregnancy carries up to 15% risk of ZIKV-associated birth defects [11–14]. This teratogenic capacity distinguishes ZIKV from other flaviviruses, yet the determinants underlying this unique pathology remain poorly understood. Determining whether this capacity extends to ZIKV’s closest relatives is critical for assessing emerging threats and developing targeted interventions.

Following its identification in 1947, ZIKV was perceived to have low pathogenic potential and little capacity to cause major outbreaks [15,16]. This perception began to shift following explosive outbreaks on Yap Island (2007) and French Polynesia (2013), both characterized by high attack rates, although pregnancy complications were not recognized at the time [17,18]. The 2015–16 Americas epidemic revealed ZIKV’s capacity to cause congenital Zika syndrome (CZS), a newly identified spectrum of fetal harm including microcephaly, neurological impairment, and fetal loss [19–23]. Given that ZIKV’s potential for fetal harm went unrecognized for decades, we posited that close genetic relatives of ZIKV might also possess this potential.

Spondweni virus (SPOV) is ZIKV’s closest known relative, sharing ∼68% nucleotide similarity and ∼75% amino acid similarity [24]. Both viruses have documented co-circulation in sub-Saharan Africa and antibodies against either virus exhibit extensive cross-reactivity with the other, complicating surveillance and diagnostic efforts [24–28]. Due to their extensive serological cross-reactivity, SPOV and ZIKV are grouped together taxonomically in the Spondweni virus serocomplex. Despite these genetic and geographic similarities, SPOV has caused only six documented human infections, most presenting as mild-to-moderate, self-limiting febrile illness [27,29–33]. These cases span from SPOV’s first identification in Nigeria in 1952 [29] to its most recent documented human case in Burkina Faso in 1979 [33]. While the limited number of reported cases presented with relatively mild symptoms, some cases have included vascular or neurological symptoms, such as epistaxis (nosebleed) [31] and photophobia [33], suggesting the possibility of more severe clinical manifestations [27].

Historically, SPOV has been detected only in sub-Saharan Africa, where it has been isolated from humans and mosquitoes [24,31,34,35]. In 2016, however, SPOV was unexpectedly detected in *Culex quinquefasciatus* mosquitoes in Haiti [36], raising the possibility that SPOV, similar to ZIKV, may have the potential to expand its geographic range beyond Africa. Given these concerns and SPOV’s close genetic relatedness to ZIKV, we sought to evaluate whether SPOV shares ZIKV’s capacity to cause fetal harm. We previously demonstrated that SPOV can cause fetal demise and placental pathology and can be vertically transmitted in a mouse pregnancy model [37]. However, mouse placental structure is labyrinthine rather than villous, as seen in humans, and human flaviviruses like DENV, ZIKV, and SPOV can only replicate in mice lacking type-I interferon responses, so it is important to evaluate their pathogenic potential during pregnancy in additional models [37–39].

To overcome these limitations, our group has used rhesus macaques to model flavivirus pathogenesis during pregnancy [40–46]. Rhesus macaques recapitulate core features of human pregnancy, including gestational timing, villous placental architecture, and immune function [47–49]. Using this model, our group has demonstrated that viral lineage and timing of infection are crucial determinants of ZIKV-associated fetal harm. In previous studies with Asian-lineage ZIKV (PRVABC59 strain; hereafter ZIKV-PR) we found infrequent fetal demise and no cases of microcephaly in pregnant rhesus macaques, though infection was associated with neurodevelopmental deficits and presence of vRNA in the maternal-fetal interface (MFI) [42,50,51]. In contrast, African-lineage ZIKV (Dak Ar 41524 strain; hereafter ZIKV-DAK) consistently caused frequent fetal demise between 12 and 20 days post-inoculation when inoculated during mid-first trimester (gestational day 30) [41]. Motivated by these observations and prior evidence that SPOV replicates in non-pregnant rhesus macaques [52], we designed an experiment to assess the comparative pathogenic potential of SPOV, ZIKV-DAK, and ZIKV-PR in pregnancy. We inoculated four pregnant rhesus macaques with SPOV and compared maternal viremia, fetal growth, and placental pathology at delivery to a previously characterized cohort of five macaques infected with ZIKV-DAK and eight macaques infected with ZIKV-PR at the same approximate gestational age [41,53]. This study tests whether SPOV, under matched conditions, produces placental infection and fetal injury comparable to ZIKV-DAK or ZIKV-PR.

## Results

### Study design

We inoculated four pregnant rhesus macaques with SPOV at approximately 30 days after the estimated date of conception (range GD 31–34; **Fig 1**). All four macaques were inoculated with 1×10^4 plaque-forming units (PFU) of SPOV (SAAr94 strain), a dose within the range thought to be delivered by mosquitoes [54]. In a previous study [41], we inoculated five pregnant rhesus macaques with 1×10^4 PFU of ZIKV (ZIKV-DAK; Dak Ar 41524 strain) at approximately GD 30. Recently [53], we also inoculated eight pregnant rhesus macaques with 1×10^4 PFU of ZIKV (ZIKV-PR; PRVABC59 strain) at approximately GD 30. In each study, we assessed maternal plasma viral load throughout the acute phase of infection and fetal growth throughout gestation until the study endpoint. In the SPOV and ZIKV-DAK studies, we also assessed fetal and placental tissue viral loads following necropsy. In the ZIKV-PR study, we assessed placental but not fetal tissues, as infants remained with their dams for developmental studies following cesarean delivery at approximately GD 160 (term in rhesus macaques is approximately 165 days) [53].

**Fig 1.**
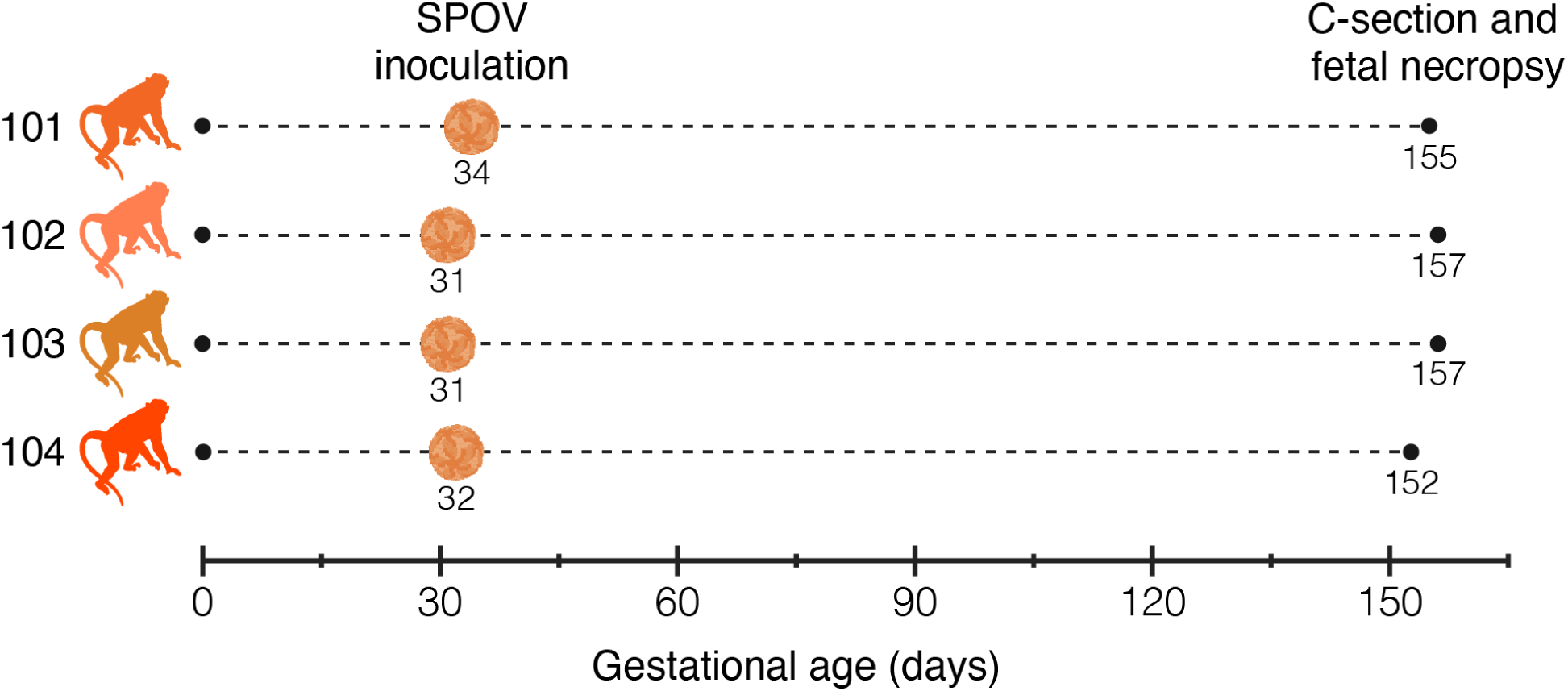
Study design. Four pregnant rhesus macaques (101–104) were inoculated subcutaneously with 1×10^4 PFU of Spondweni virus (SPOV) between gestational days (GD) 31–34. Cesarean-section delivery, euthanasia, and fetal necropsy were performed near term (GD 152–157) in all animals; dams survived the procedure.

### Pregnant macaques support SPOV infection with occasional prolonged shedding

All SPOV-inoculated macaques became productively infected, reaching a median peak maternal plasma viral loads of 2.51×10^4 copies/mL (range: 1.38×10^4 – 3.85×10^5 copies/mL) approximately six days post-inoculation (**Fig 2A**). Plasma viremia persisted for a median duration of 16.5 days (range: 10–31 days). We define prolonged viremia as ≥21 days of detectable plasma vRNA on serial sampling, because in non-pregnant rhesus macaques challenged with 1×10^4 PFU of SPOV, ZIKV-DAK, or ZIKV-PR, plasma viremia typically resolves within ten days [52]. By this definition, macaques 102 and 104 exhibited prolonged viremia (31 and 23 days, respectively), whereas macaques 101 and 103 were viremic for ten days each. All macaques were serologically naïve to SPOV prior to inoculation and seroconverted by 30 days post-inoculation (**S1 Fig**).

**Fig 2.**
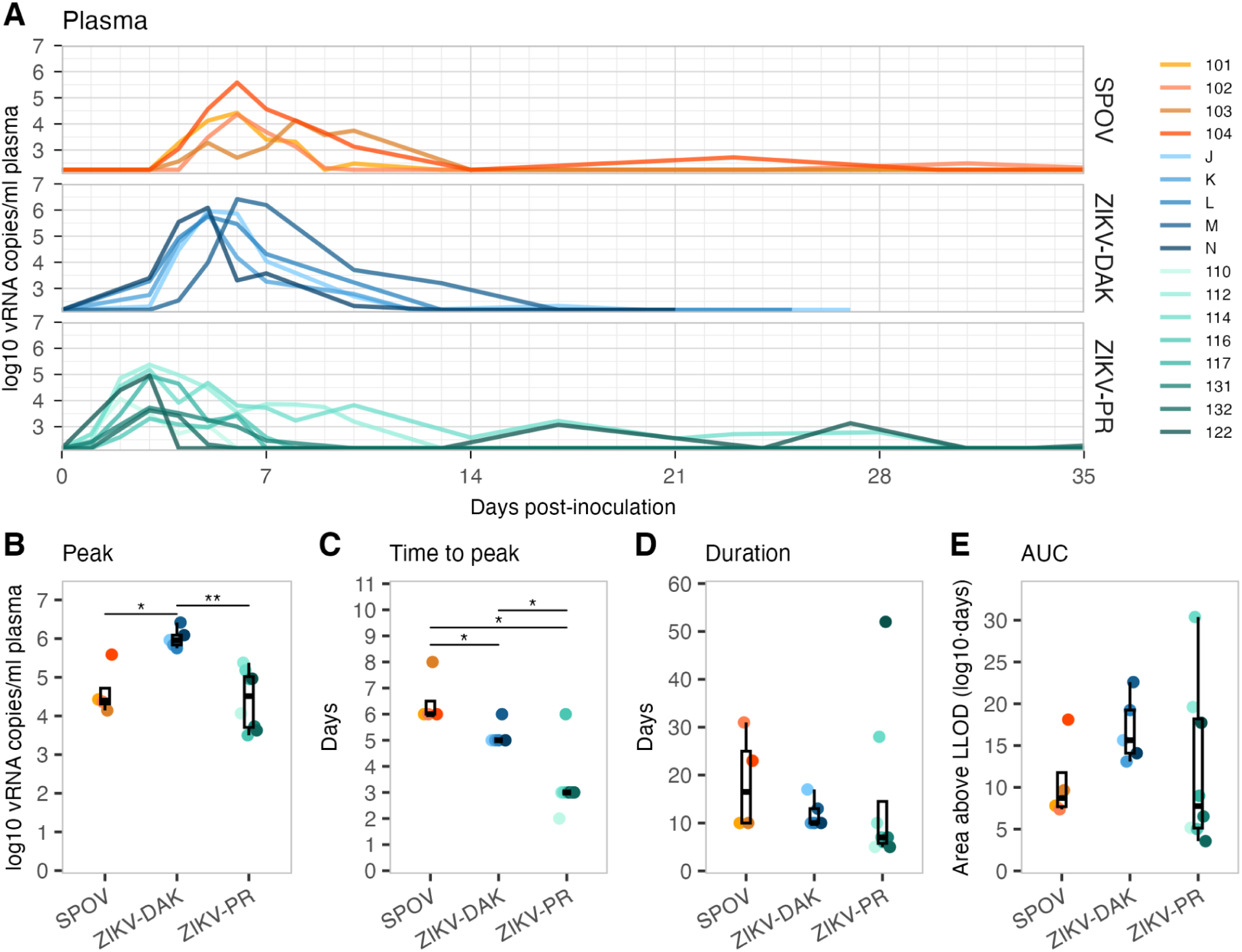
Plasma viremia kinetics in pregnant macaques after Spondweni or Zika virus inoculation. **(A)** Longitudinal plasma viral RNA loads in pregnant macaques inoculated with Spondweni virus (SPOV; orange), Zika virus Dak Ar 41524 strain (ZIKV-DAK; blue), or Zika virus PRVABC59 strain (ZIKV-PR; green). Viral loads are shown as log10 RNA copies per mL of plasma over days post-inoculation. **(B–E)** Group comparisons of **(B)** peak viral load, **(C)** time to peak viremia (days), **(D)** duration of viremia (days above the assay lower limit of detection, LLOD), and **(E)** area under the curve (AUC) defined as area above the LLOD (units: log10·days).

In comparison, ZIKV-DAK-infected macaques had higher peak maternal plasma viral loads than those infected with SPOV (9.12×10^5 copies/mL; BH-adjusted *p* = 0.0238), whereas peak viremia for SPOV was not significantly different from ZIKV-PR (*p* = 0.570; **Fig 2B**). In the SPOV group, viral loads peaked approximately six days post-inoculation (**Fig 2C**), significantly later than both ZIKV-DAK (five days; *p* = 0.0397) and ZIKV-PR (three days; *p* = 0.0242). The duration of viremia (**Fig 2D**) did not differ significantly between SPOV and ZIKV-DAK (ten days; *p* = 0.524) or SPOV and ZIKV-PR (seven days; *p* = 0.345). To quantify the overall burden of viral replication, we calculated the area under the curve (AUC) for each animal, excluding values at or below the lower limit of detection. AUC did not differ significantly across pairs, although the ZIKV-DAK group had an approximately two-fold greater AUC compared to the SPOV group (*p* = 0.333; **Fig 2E**). Collectively, these findings demonstrate that SPOV-inoculated pregnant rhesus macaques exhibited later peak viremia than animals infected with either of the ZIKV strains.

While the total burden of virus replication trended lower for SPOV compared to ZIKV-DAK, this difference was not significant.

Previously, we detected ZIKV RNA in urine from all five ZIKV-DAK-inoculated macaques [41], with one animal reaching a peak urine viral load of nearly 48,000 copies/mL. In this study, we detected SPOV RNA in urine from two SPOV-inoculated macaques (101 and 104; **S2 Fig**). Macaque 101 had detectable viral RNA (vRNA) at 118 copies/mL five days post-inoculation, increasing to 1,016 copies/mL by day ten. Macaque 104 had detectable vRNA at 159 copies/mL nine days post-inoculation. These results suggest that SPOV, similar to ZIKV-DAK, can be detected in urine but appears to exhibit more limited urinary shedding in this rhesus macaque model.

Points represent individual macaques (SPOV *n* = 4; ZIKV-DAK *n* = 5; ZIKV-PR *n* = 8). Horizontal brackets in **B** and **C** mark pairwise two-tailed Wilcoxon rank-sum tests. Stars denote Benjamini-Hochberg-adjusted p-values within each metric (* < 0.05, ** < 0.01, *** < 0.001); non-significant p-values are not shown. ZIKV-DAK-inoculated macaques were previously reported by Rosinski and Raasch *et al*. [41]. ZIKV-PR-inoculated macaques were previously reported by Krabbe *et al*. [53].

### Prenatal ultrasound reveals no consistent evidence of SPOV-associated intrauterine growth restriction

All pregnancies in SPOV-inoculated macaques progressed to the study endpoint (GD 152–157; term = 165 ± 10 days) without fetal demise, premature birth, or gross anatomical abnormalities. This finding contrasts with our prior study of ZIKV-DAK-inoculated macaques inoculated at the same point in the first trimester; each of those pregnancies ended in fetal demise by ∼20 days post-inoculation [41]. This suggests that SPOV has reduced pathogenic potential compared to ZIKV-DAK, but the absence of overt adverse outcomes does not preclude more subtle effects on fetal development. To monitor fetal growth, we conducted biweekly ultrasounds from early gestation through to near-term (gestational day 155), measuring fetal biparietal diameter (BPD), head circumference (HC), abdominal circumference (AC), and femur length (FL). Growth trajectories appeared broadly consistent and uninterrupted across gestation (**Fig 3A**). To contextualize these trajectories, we compared our measurements to age-matched normative data from a large California National Primate Research Center (CaNPRC) cohort of rhesus macaques (*n* = 55) [55,56]. We then calculated Z-scores to quantify the extent to which each measurement deviated from the CaNPRC mean (**Fig 3B**). SPOV-exposed fetuses showed statistically significant deviations from expected mean Z-score values for all measurements (*p* < 0.05), but their growth trajectories (slope of the Z-score curves) were not significantly different from the normative data (**Fig 3B**). All SPOV-exposed fetuses had HC that measured below the CaNPRC mean throughout gestation (p < 0.001; **Fig 3A**), but differences were small. In human fetal medicine, the threshold for abnormal size is defined as a Z-score ≥2, and Z-scores for SPOV-exposed fetuses never reached this threshold [57,58]. The deviation in HC was most pronounced after 15 weeks of gestation, where fetus 003 (from macaque 103) had an HC Z-score of -1.98 at 20 weeks of gestation, just under the abnormality threshold (**S3A Fig**).

**Fig 3.**
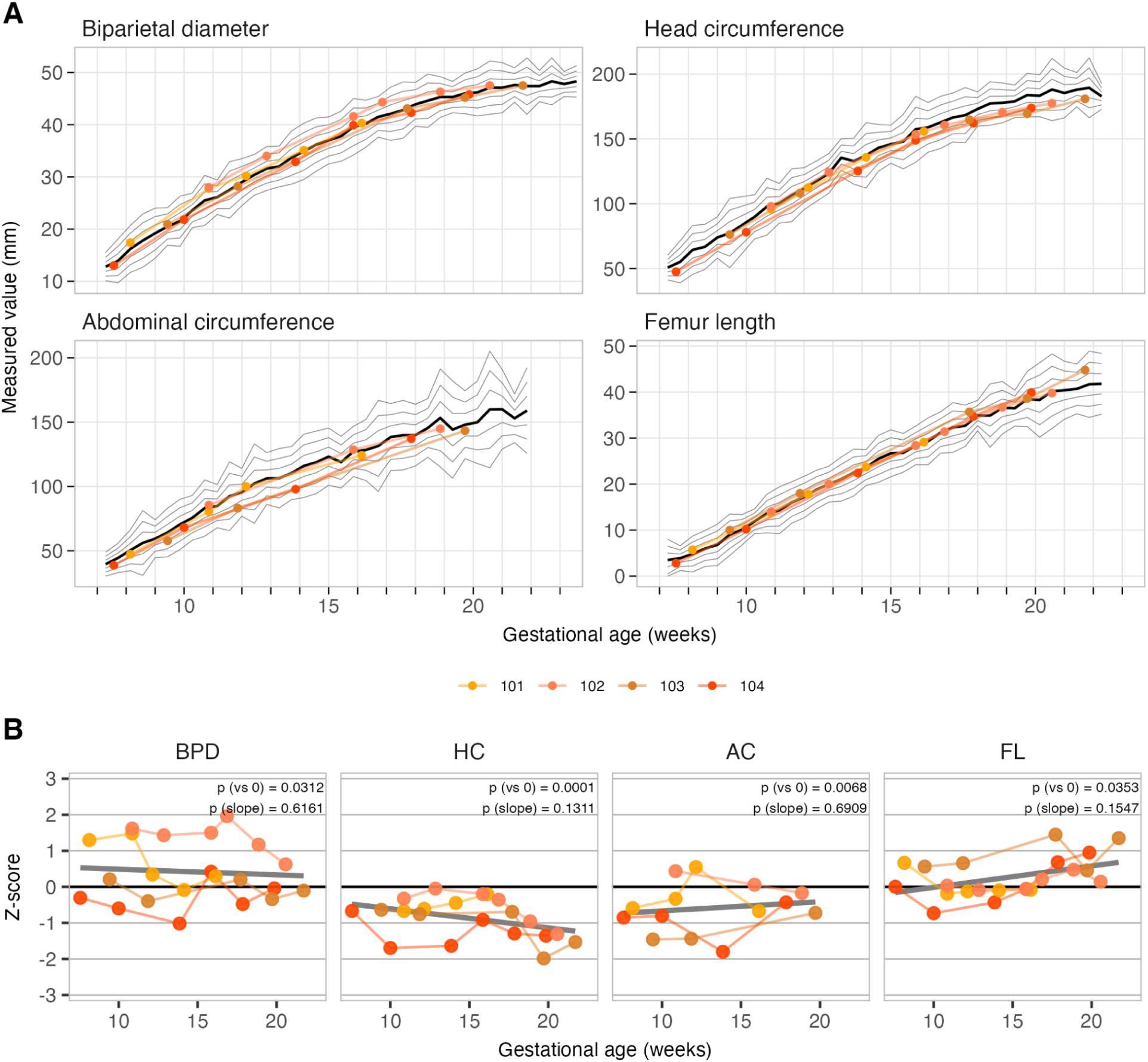
Fetal growth trajectories in Spondweni virus-inoculated macaque pregnancies. **(A)** Fetal growth measurements from the macaques inoculated with Spondweni virus (SPOV): biparietal diameter (BPD), head circumference (HC), abdominal circumference (AC), and femur length (FL). Points denote individual macaques (*n* = 4), with lines connecting repeated measurements over gestational age (weeks). Normative reference curves are shown as a black line (mean) with flanking grey lines representing ±1, ±2, and ±3 standard deviations at each gestational age, derived from rhesus macaque reference cohorts (*n* = 55) [55,56]. **(B)** Z-scores for SPOV-inoculated macaques at each time point for the same four metrics. Values reflect the number of standard deviations each measurement differed from the gestational-age-matched normative mean. A linear regression line (dark grey) is overlaid in each panel. *p*-values indicate one-sample t-tests (vs 0) and a linear mixed-effects model (slope). Horizontal grey lines indicate ±1, ±2, and ±3 Z-score thresholds, corresponding to the mean-flanking grey lines in **A**.

BPD was not as markedly reduced (**Fig 3A**) and remained closer to the normative mean, suggesting that the observed reduction in HC may reflect differences in cranial shape or growth pattern rather than a global restriction in fetal head size. AC was also statistically significantly smaller in SPOV-exposed fetuses than in the normative data (*p* < .01), but FL was significantly elevated (*p* < 0.05; **Fig 3B**). Together our fetal growth data suggest that SPOV-exposed fetuses in our study were somewhat smaller than fetuses in the normative dataset, but that their growth followed a similar trajectory. It appears likely that size differences relative to the normative cohort were statistically, but not biologically, significant. Comparison of our SPOV-exposed fetuses with the growth trajectories of other rhesus macaques studied at the Wisconsin National Primate Research Center (WNPRC), including mock-infected (*n* = 4), ZIKV-DAK (GD 30; *n* = 4), ZIKV-PR (GD 30; *n* = 8) and ZIKV-PR cohorts (GD 45; *n* = 4), supports this conclusion: we observed no apparent differences in growth trajectories between our SPOV-exposed animals and these groups (**S3B Fig**) [42]. Thus, the HC and AC values of WNPRC fetuses may consistently fall below CaNPRC normative curves, regardless of viral exposure. Together, our observations indicate that SPOV does not cause an apparent phenotype of fetal growth restriction in rhesus macaques.

### SPOV RNA is present at low levels in maternal and fetal tissues upon delivery

Despite the lack of overt fetal growth restriction, we next asked whether SPOV RNA was detectable in maternal, fetal, or maternal-fetal interface (MFI) tissues at delivery. SPOV RNA was sporadically detected at very low levels in placental tissues and the maternal spleen at the time of fetectomy (**Fig 4**). Specifically, vRNA was identified in the spleens of SPOV-infected macaques 101 and 102, with macaque 101 testing positive in one of two technical replicates (0 and 7 copies/mg) and macaque 102 testing positive in both wells (34 and 40 copies/mg) (**Fig 4**; **Table 1**). Additionally, we detected 10 copies/mg in the decidua basalis of macaque 101. In macaque 104, we detected SPOV RNA in various samples of the decidua basalis (1–4 copies/mg across samples), placental parenchyma tissue (4–8 copies/mg), and chorionic plate (5–42 copies/mg; **Fig 4**). This pattern of low-level tissue vRNA burden resembles our findings with ZIKV-PR (mesenteric lymph node and MFI: decidua basalis and placental parenchyma). In contrast, we previously detected ZIKV-DAK RNA in maternal organs (liver, spleen, mesenteric lymph node) and multiple MFI compartments, frequently at higher levels. Because tissues from animals infected with ZIKV-DAK were sampled at the time of fetal demise (within 20 days of inoculation), whereas tissues from SPOV- and ZIKV-PR-infected pregnancies were sampled much later, over 100 days post-inoculation, we cannot determine from these data whether higher ZIKV-DAK tissue burdens reflect inherently greater replication in the MFI or earlier sampling for ZIKV-DAK during gestation. Per-animal tissue vRNA loads are summarized in **Table 1** for SPOV-inoculated macaques. SPOV RNA was undetectable across 60 fetal anatomical compartments, including CNS, ocular, cardiopulmonary, lymphoid, gastrointestinal, reproductive, and endocrine (**S1 Table**). Because we previously found evidence of ocular pathology in fetuses from ZIKV-infected dams [59], eyes of four SPOV-exposed fetuses were analyzed histopathologically via light microscopy, but no abnormalities (developmental, inflammatory or otherwise) were identified (data not shown).

**Fig 4.**
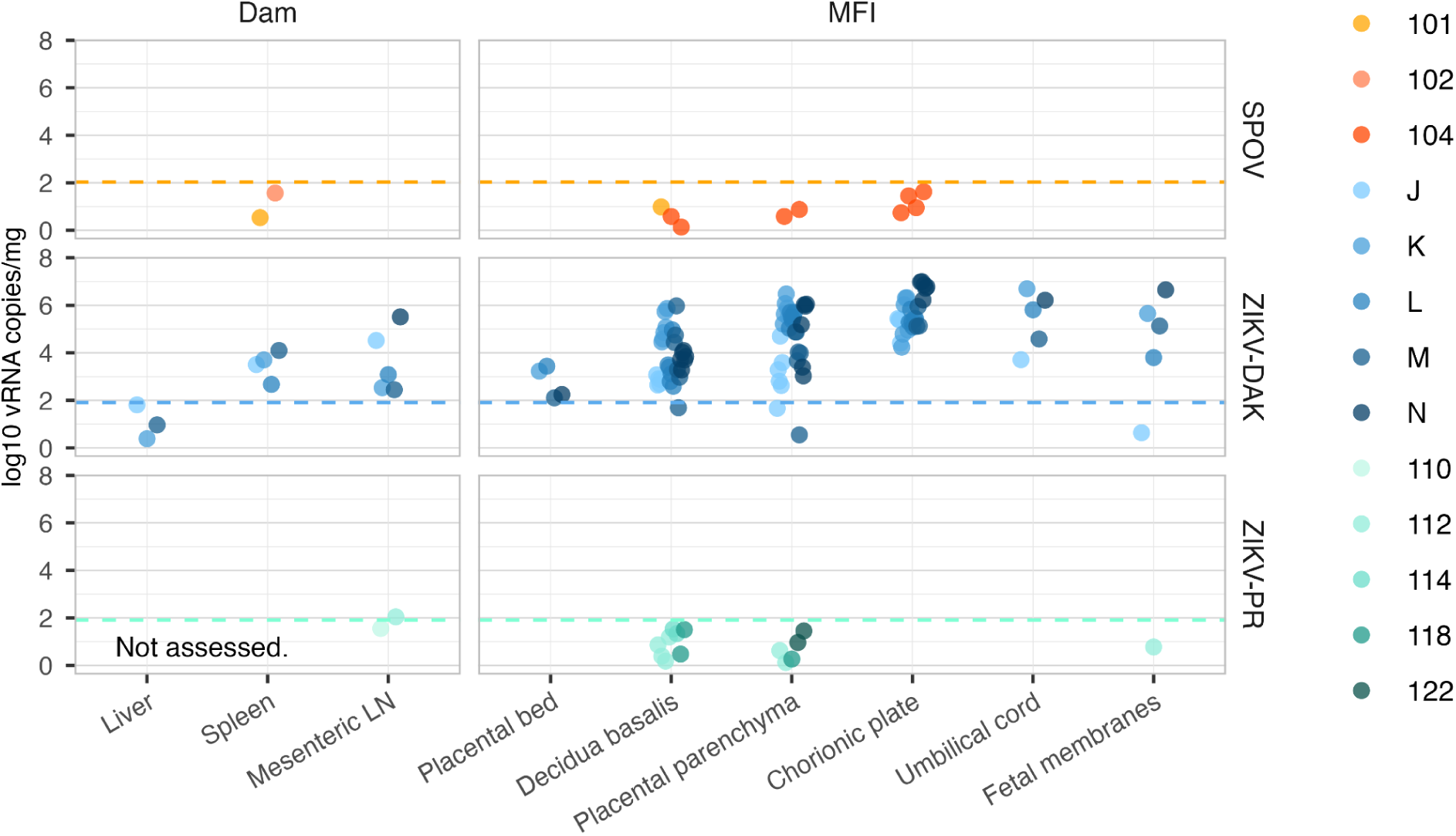
Tissue viral RNA at delivery in dams and the maternal–fetal interface. Viral RNA levels (log10 copies per mg tissue) measured at delivery from pregnant macaques inoculated with Spondweni virus (SPOV; orange), Zika virus Dak Ar 41524 strain (ZIKV-DAK; blue), or Zika virus PRVABC59 strain (ZIKV-PR; green) earlier in gestation. Tissues are grouped into maternal (Dam) and maternal-fetal interface (MFI) compartments. Each point is a single sample from one animal (SPOV *n* = 4; ZIKV-DAK *n* = 5; ZIKV-PR *n* = 8). Only samples with detectable viral RNA are shown; dam 103 and her fetus had no detectable vRNA. The horizontal dashed line indicates the assay’s lower limit of detection (LLOD = 108 copies/mg for SPOV and 81 copies/mg for ZIKV), defined as the concentration detected ≥95% of the time in validation experiments (see Methods). Values plotted below this threshold reflect occasional true-positive detections but are not reliably quantifiable. ZIKV-DAK-inoculated macaques were previously reported by Rosinski and Raasch *et al*. [41]. ZIKV-PR-inoculated macaques were previously reported by Krabbe *et al*. [53].

**Table 1.** Virological and pathological findings.

| ID | Maternal spleen vRNA<sup>1</sup> | MFI vRNA<sup>1</sup> | Fetal vRNA<sup>1</sup> | Chronic chorionic villitis<sup>2</sup> | Maternal vascular malperfusion | Villous ischemic injury<sup>2</sup> | Trans-mural ischemia<sup>2</sup> |
| --- | --- | --- | --- | --- | --- | --- | --- |
| 101 | ±<sup>3</sup> | + | – | + | – | + | + |
| 102 | + | – | – | + | + | + | + |
| 103 | – | – | – | + | + | + | + |
| 104 | – | +<sup>4</sup> | – | + | + | + | + |

<sup>1</sup>Detection of Spondweni vRNA by qRT-PCR (duplicate wells per tissue). In this table + = both wells amplified (2/2); ± = equivocal (1/2); – = neither well amplified (0/2). All detections were below the assay lower limit of detection (LLOD = 108 copies/mg tissue) and are not reliably quantifiable. All fetal tissues (60 compartments) were negative (see **S1 Table** for the full list of tissues tested). Abbreviations: vRNA, viral RNA; MFI, maternal-fetal interface.
<sup>2</sup>Pathology scoring on H&E. (+) present; (–) not observed. Chronic chorionic villitis refers to mononuclear/lymphohistiocytic chorionic villitis. Transmural ischemia was defined as any plate-to-plate area of ischemic necrosis present on any full-thickness section from either placental disc. See **S2 Table** for a summary of placental histopathological findings.
<sup>3</sup>101 maternal spleen: one of two wells amplified (0 and 7 copies/mg).
<sup>4</sup>104 MFI: one placental parenchyma sample was equivocal, and a second parenchyma sample tested positive in both wells (7 and 8 copies/mg). Decidua basalis (two samples) and chorionic plate (four samples) tested positive but below the LLOD in both wells.

### SPOV exposure induces placental inflammation and vascular pathology

We reasoned that placental histopathology might reveal signs of earlier infection or vascular injury during acute maternal viremia, even in the absence of gross fetal abnormalities or detectable SPOV RNA in fetal tissues at delivery. To investigate this possibility, we examined H&E-stained full-width and full-thickness central sections (“center cuts”) from the midpoint of each placental disc together with representative sections from each cotyledon collected at cesarean section. Macaques typically have bidiscoid placentas, and both placental discs were sampled per pregnancy. In the placentas from 3 of 4 SPOV-inoculated dams (102, 103, and 104), we observed persistently muscularized arteries in the decidua basalis (**Fig 5A**), suggestive of impaired maternal spiral-artery remodeling, a hallmark of maternal vascular malperfusion (MVM; for a normal, remodeled maternal spiral artery, see the bottom of **S4A Fig**, showing a large basal-plate artery with a wide lumen and thin vessel wall). These findings were accompanied by dense clusters of cells with segmented (multilobed) nuclei, morphologically consistent with neutrophilic infiltration, in the decidua and small proximal vessels (**Fig 5B**), indicating acute (neutrophilic) deciduitis and vasculitis. Larger decidual vessels, in contrast, exhibited predominantly mononuclear infiltrates composed of small round lymphocyte-like cells and evidence of prior hemorrhage, including interstitial hemosiderin deposition and hemosiderophages. and fibroblast disintegration (**Fig 5C–D**). Per-animal summaries are provided in **S2 Table**. These findings are consistent with mononuclear inflammation of the larger decidual vessels. In aggregate, these findings suggest that SPOV infection can induce placental inflammation and vascular pathology that may impair maternal-fetal blood exchange, even in the absence of overt fetal harm.

**Fig 5.**
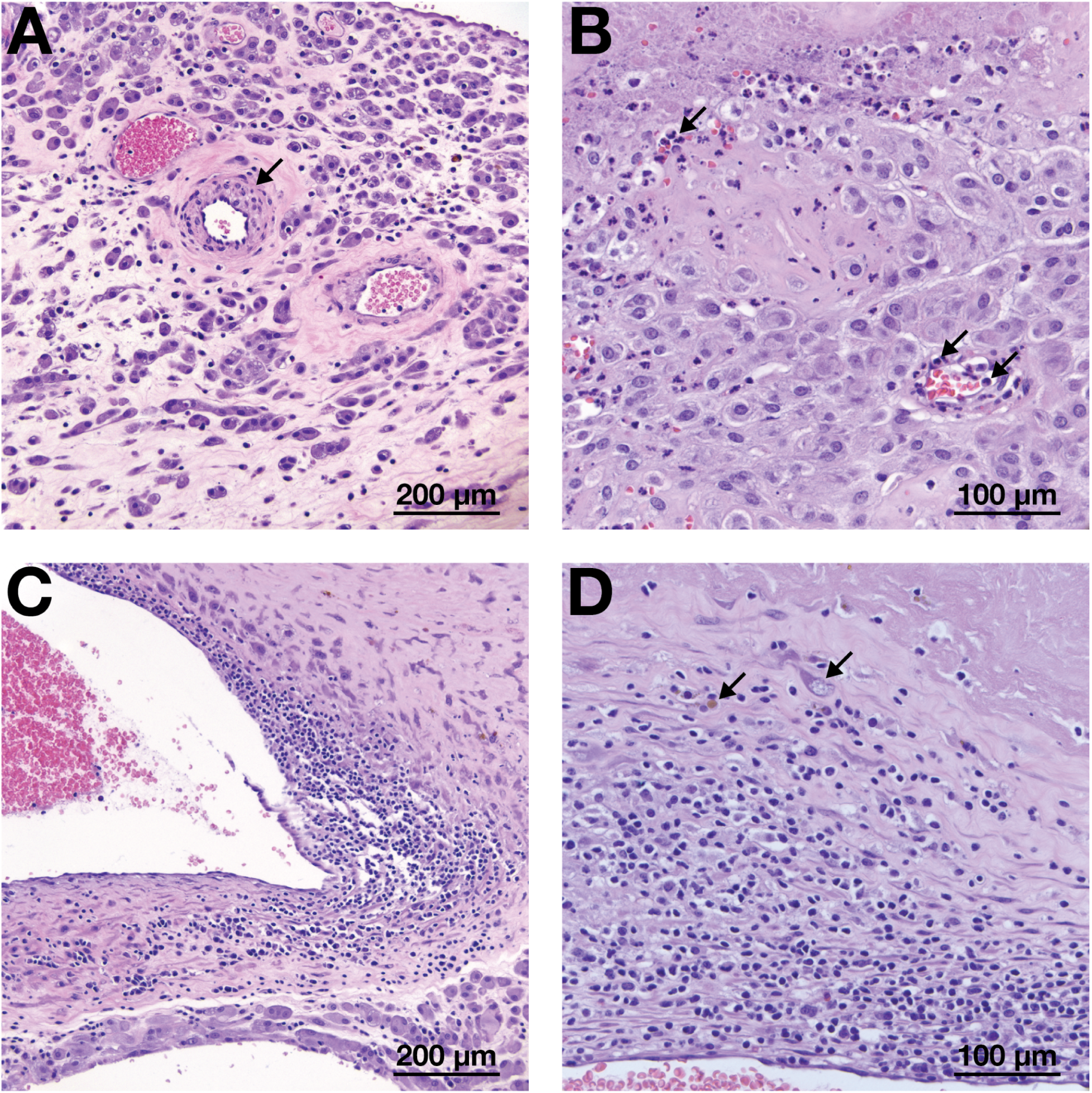
Representative images of placental blood vessel histopathology. **(A)** Persistently muscularized artery (black arrow) in the decidua basalis from macaque 103. **(B)** Neutrophilic deciduitis (top arrow) and vasculitis (bottom arrows) of a small vessel in the decidua parietalis from macaque 101. The bottom arrows show neutrophils in the lumen (right-most arrow) and epithelium (left-most arrow) of the vessel. **(C)** Mononuclear/lymphohistiocytic vasculitis of a large artery within the basal plate of macaque 102. **(D)** Mononuclear/lymphohistiocytic vasculitis of a large basal plate vessel from macaque 102, with the left-most arrow showing a hemosiderin deposit and the right-most arrow showing a fibroblast with a disintegrating nucleus.

We next evaluated the histopathology of the chorionic villi, which mediate the oxygenation of fetal blood. All four SPOV-exposed macaques exhibited evidence of villous injury, although the severity and histological features varied. A representative image of normal villous architecture (**S4B Fig**) shows branching villi without evident inflammation. Macaques 102, 103, and 104 showed multifocal mononuclear/lymphohistiocytic villitis with associated perivillous fibrin deposition, mineralization, and ischemic necrosis. Among these, macaque 103 exhibited the most severe villous pathology, with widespread necrosis and collapse of villous architecture (**Fig 6A**). Macaque 102 displayed moderate villitis and hemorrhage, along with diffuse perivillous fibrin and mineral accumulation within the basal plate and adjacent villi (**Fig 6B**). Macaque 104 showed mononuclear/lymphohistiocytic villitis accompanied by small plasma cell aggregates consistent with chronic deciduitis (**Fig 6C**). Most strikingly, macaque 104 also had extensive transmural placental ischemic necrosis and loss of parenchymal villi that caused the chorionic plate to collapse into a distinctive “V” configuration (**S5 Fig**). In contrast, macaque 101 had relatively mild mononuclear/lymphohistiocytic chorionic villitis with focal villous ischemic injury, including small areas of necrosis, thrombosis, and mineralization (**Fig 6D**). Across animals, MVM was present in 3/4 and villous ischemic injury in 4/4, and transmural ischemia was present in all four placentas by our definition (any plate-to-plate region of ischemic coagulative necrosis), with the most striking example observed in macaque 104 (**Table 1**). The spectrum of villous injury by animals is summarized in **S2 Table**. Together, these findings demonstrate that SPOV exposure can result in diverse villous pathology ranging from focal inflammation to extensive multifocal ischemia, highlighting that significant placental pathology may occur without translating to overt fetal harm. Two of four infants did have minimal to mild multifocal neutrophilic inflammation involving the liver (fetuses 002 and 003 from macaques 102 and 103, respectively), spleen (fetus 003), salivary gland (fetus 002), and axillary lymph node (fetus 002). No other organs, including the eyes and brain, had significant histologic changes in any of the infants.

**Fig 6.**
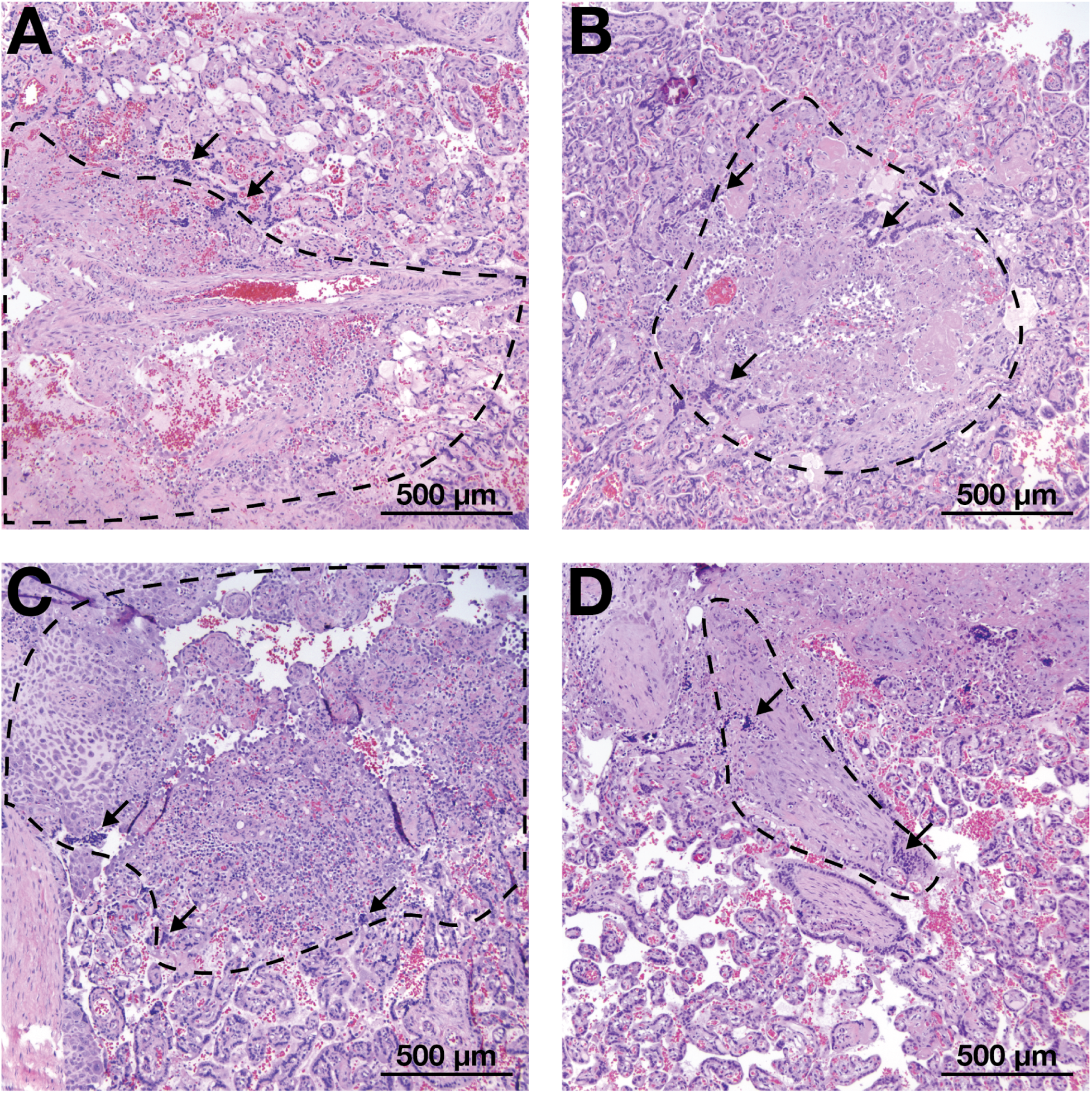
Representative images of placental chorionic villi histopathology. (A) Mononuclear/lymphohistiocytic chorionic villitis in Macaques 103, **(B)** 102, **(C)** 104, and **(D)** 101. In each image, representative clusters of lymphohistiocytic infiltration are denoted by black arrows, and the loss of villous architecture is circumscribed by a dashed black line.

## Discussion

ZIKV and SPOV are close relatives and the two known members of the Spondweni virus serocomplex. It is currently unknown whether SPOV has the potential to cause fetal harm. This study provides the first assessment of SPOV infection during pregnancy in a translational rhesus macaque model. We demonstrate that SPOV induces placental pathology, yet does not cause fetal demise or gross fetal abnormalities when fetuses were collected just prior to term. Nevertheless, SPOV was associated with placental pathology in all macaques and is likely able to infect the placenta.

We found very low levels of SPOV RNA, below the assay’s lower limit of detection, in several tissue specimens from 3 of 4 dams and/or fetuses. We believe these findings represent bona fide detection of SPOV genomic material for several reasons. RT-PCR assays can detect vRNA at lower concentrations, but below this limit we expect detection of true positives to become increasingly sporadic. Values below the LLOD are therefore detectable but not reliably quantifiable, limiting our ability to draw conclusions about prolonged viral replication at these sites. Extraction-negative and no-template controls were uniformly negative, and positive controls were uniformly positive on every instrument run. Presence of SPOV RNA is also consistent with findings of pathological changes in the placenta of some SPOV-infected dams. Together, our data therefore argue against cross-contamination and suggest true SPOV RNA presence at very low levels in maternal and fetal tissues ∼120 days after inoculation.

Placental pathology in all SPOV-inoculated macaques and low, but detectable, levels of vRNA at term (∼120 days post-inoculation) in maternal-fetal interface tissues of *n* = 2/4 macaques suggest that SPOV is able to induce placental injury that can persist even in the absence of detectable fetal infection (**Table 1**). This is similar to what we observed with the Asian-lineage ZIKV strain PRVABC59 but direct comparisons of placental viral burdens with ZIKV-DAK are complicated by sampling at different timepoints: ZIKV-DAK tissues were collected at the time of fetal demise (within 20 days of inoculation), when some dams remained viremic; SPOV and ZIKV-PR tissues were collected more than 100 days after inoculation, long after maternal viremia had resolved. Findings of mononuclear/lymphohistiocytic chorionic villitis and maternal vascular malperfusion suggest that SPOV induced placental damage early in pregnancy, triggering lasting inflammatory damage and impaired spiral artery remodeling.

Further, SPOV induced neutrophil-predominant inflammation in the decidua parietalis of macaque 101, which also contained low levels of vRNA in the decidua basalis, together suggesting that SPOV is capable of prolonged viral persistence and sustained inflammatory responses in both the placenta and the fetal membranes. Last, SPOV can induce severe placental pathology: macaque 104, which contained low vRNA in the decidua, parenchyma, and chorionic plate, developed significant regions of ischemia and villous necrosis **S5 Fig**). These observations indicate that, despite considerable pathology and vRNA presence upon delivery, placental structure and function remained resilient enough to prevent any detectable vertical transmission of SPOV or notable injury to the fetus.

Gestational timing of viral inoculation profoundly affects ZIKV congenital outcomes in non-human primate models. In previous studies, inoculation later in the first trimester (around GD 45) with African-lineage ZIKV-DAK or Puerto Rican ZIKV-PRVABC59 was not consistently associated with fetal demise or marked fetal growth restriction, and in some cohorts no ZIKV RNA was detected in infants at delivery. [42,51,60,61]. Vertical transmission can occur at this gestational age: inoculation in the mid-first trimester (GD 31-38) with a French Polynesian strain of ZIKV (ZIKV-FP) caused congenital infection with fetal ocular injury [46]. In contrast, when inoculated at GD ∼30, ZIKV-DAK caused uniform fetal demise and vertical transmission 12-20 days after inoculation [41]. Notably, infectious ZIKV-DAK has been recovered from amniotic fluid within ∼2 weeks of inoculation at GD ∼45 in at least one pregnancy, indicating that African-lineage ZIKV can also cause detectable fetal infection, particularly when the fetal compartment is sampled soon after infection [62]. Similarly, the absence of severe fetal outcomes in our SPOV inoculations at GD ∼30 does not exclude the possibility of SPOV posing a congenital infection risk in humans.

A variety of viral and host factors likely contribute to these differences in infection outcomes. The stark contrast in fetal survival rates for dams inoculated with ZIKV-DAK at GD 30 vs. GD 45 suggests that there is a critical window during the mid-first trimester when the fetus is particularly susceptible to infection, perhaps because the MFI is still undergoing rapid trophoblast invasion and spiral-artery remodeling at GD 30, while these processes are more advanced by GD 45 [41]. Inoculum dose likely also plays a role. All infected dams whose viremia data are summarized in this manuscript were inoculated with 1×10^4 PFU virus, an amount in the middle of the range thought to be delivered by mosquitoes [54]. Notably, we previously showed that 1×10^8 PFU ZIKV-DAK inoculated at GD ∼45 led to fetal infection, indicating that high doses can overcome barriers to vertical transmission that develop over the course of the first trimester [40]. However, strain differences are prominent: inoculation with 1×10^4 PFU of ZIKV-FP at GD 31–38 resulted in vertical transmission with fetal ocular injury [46]. Across experiments that maintained the same inoculum dose (1×10^4 PFU), inoculation route, gestational timing (GD ∼30), and animal care conditions, outcomes diverged by virus: ZIKV-DAK produced uniform fetal demise with vertical transmission [41], whereas in the present study, SPOV produced neither fetal demise nor detectable vertical transmission despite consistent placental pathologies. Indeed, SPOV infection more closely resembled our previous observations of ZIKV-PR infection at GD 30, in which very low levels of vRNA were detected in MFI tissues near the time of delivery. Interestingly, the duration of infection during pregnancy appeared more variable for ZIKV-PR—with viremia lasting over 50 days in one dam—than we observed for SPOV, which was cleared from plasma within 30 days by all four dams in this study (**Fig 2**). Taken together, our studies therefore indicate that viruses within the Spondweni virus serocomplex likely possess a wide range of pathogenic potential during gestational infection. Improving risk assessments for these viruses will require further studies to identify specific viral and host determinants of pathogenic outcomes.

Several limitations constrain our findings. Our data, from a small cohort (*n* = 4) with one SPOV strain, dose, and inoculation at a single gestational time point, suggest that SPOV can damage the placenta without demonstrable fetal infection or gross abnormalities observable at term. Because fetal tissues were uniformly negative for SPOV vRNA while placentas showed consistent pathology (and occasional low-level MFI vRNA), our findings are most consistent with placenta-focused infection and injury rather than direct infection of the fetus. Whether such injury manifests in postpartum impacts such as neurodevelopmental effects is unknown and would require long-term follow-up. Further, our study design, which terminated pregnancies at near-term rather than during acute infection, may have reduced our ability to detect vertical transmission or characterize acute placental responses. To understand the placental response and vRNA distribution during acute maternal infection, future studies should include necropsies at earlier post-inoculation timepoints [63]. Additionally, longer-term studies should assess comprehensive developmental outcomes, which are critical to detect the subtle sequelae that we know ZIKV is capable of causing. The clinical challenge lies in recognizing that the absence of gross fetal abnormalities does not indicate the absence of risk; moreover, while cohort data exist for Asian-lineage ZIKV, comparable human data are lacking for SPOV and for African-lineage ZIKV, so we are unable to determine whether SPOV poses a different risk to human pregnancies.

The contrast between the consistent placental pathologies of SPOV and the direct fetal effects of ZIKV-DAK suggests that distinct or sequential pathways may underlie placental and fetal outcomes. This positions SPOV as a valuable comparative model for identifying the viral and host factors governing adverse pregnancy outcomes. Beyond its utility as a comparator, SPOV’s impact on the placenta raises important clinical concerns. While we observed no evidence of direct fetal harm in this cohort, the placental pathologies suggest the potential for indirect developmental effects through placental dysfunction [64]. These outcomes were not assessed in this study design but remain biologically plausible, given similar findings in ZIKV-exposed infants with developmental impairment [65,66]. These insights underscore the need for mechanistic studies of flavivirus-induced placental and fetal injury, as well as potential long-term sequelae. The severity of placental pathology we observed here warrants sustained surveillance and research in the context of expanding mosquito vector ranges and regions with ecological conditions favorable for flavivirus emergence.

## Materials and Methods

### Ethical approval

This study was approved by the University of Wisconsin College of Letters and Sciences and the Vice Chancellor for Research and Graduate Education Centers Institutional Animal Care and Use Committee (protocol numbers G006139 and G006256). The University of Wisconsin–Madison Institutional Biosafety Committee approved this work under protocol numbers B00000117 and B00000182.

### Care and use of rhesus macaques

All rhesus macaques involved in this study were housed and cared for by the Wisconsin National Primate Research Center (WNPRC) staff, as previously described by our team [40–44,46,52,67]. Care practices strictly adhered to the guidelines outlined in the Animal Welfare Act, the National Research Council’s Guide for the Care and Use of Laboratory Animals, and the Weatherall report [68].

As previously described [40–44,46,52,67], the macaques’ diets included a variety of fruits, vegetables, nuts, cereals, seed mixtures, yogurt, peanut butter, popcorn, and marshmallows. In addition to dietary enrichment, structural enrichment (e.g., climbing structures) and manipulanda (e.g., toys) were provided to promote psychological well-being. Trained animal care staff evaluated all study animals twice daily for signs of pain, distress, or illness by observing appetite, stool quality, activity level, and overall physical condition.

Macaques displaying abnormal clinical signs received appropriate care from attending veterinarians. Prior to all experimental procedures, including virus inoculation, blood collection, and physical examinations, animals were anesthetized with an intramuscular dose of ketamine (10 mg/kg) and closely monitored until fully recovered.

### Study design

In this study, we infected four pregnant rhesus macaques (*Macaca mulatta*) with SPOV and compared their viral loads and fetal development with five macaques infected with African-lineage ZIKV and eight macaques infected with Asian-lineage ZIKV. Animals that met study criteria (no prior flavivirus exposure history or major surgery) were confirmed pregnant by ultrasound and subcutaneously inoculated during the first trimester (approximately gestational day 30) with 1×10^4 plaque-forming units of either SPOV (SAAr94), African-lineage ZIKV (Dak Ar 41524), or Asian-lineage ZIKV (PRVABC59). All macaques used in the study were screened to confirm that they were not infected with the following viruses at the time of inoculation: Macacine herpesvirus 1, simian retrovirus type D (SRV), simian T-lymphotropic virus type 1 (STLV), and simian immunodeficiency virus (SIV). Pan collection of urine via removable cage bottoms was performed at all sampling time points. Blood samples were obtained from the femoral or saphenous vein using a Vacutainer system or needle and syringe. Blood was collected on all dams prior to virus exposure (0 days post-inoculation; 0 dpi), daily from 1-10 dpi, twice weekly until two sequentially negative maternal plasma vRNA loads, then once weekly until study completion. Plasma and urine samples were collected at all time points; serum samples were collected at 0, 2, 4, 7, 10, 14, 21, and 30 dpi; peripheral blood mononuclear cells (PBMCs) were collected on 0, 7, 14, 21, and 30 dpi. Pregnancies were monitored throughout gestation using ultrasound (described below) to assess fetal viability and ensure the well-being of both the dam and the fetus. Fetal outcomes were assessed upon cesarean-section delivery near GD 155, approximately 10 days prior to full term.

### Viral inoculations

The SPOV isolate (SAAr94) was derived from a *Mansonia uniformis* mosquito before five passages in unknown conditions, one passage in Vero cells, and two passages in C6/36 (*Aedes albopictus*) cells. Contemporary isolates of SPONV do not exist; thus, we used the only available low-passage isolate. Our SPOV challenge stock is 98.8% nucleotide identical to the SPOV genome recovered from mosquitoes in Haiti (GenBank: MG182017; SRA experiment accession: SRX7395146), but we acknowledge that although the sequences are almost identical, the slight difference could result in important phenotypic impacts. The ZIKV-DAK isolate (Zika virus/Aedes africanus-tc/SEN/1984/41524-DAK) was derived from *Ae. africanus* mosquitoes before two passages in *Ae. pseudocutellaris*, two passages in Vero cells, and two passages in C6/36 cells (BEI Resources; Manassas, Virginia; NR-50338). The ZIKV-PR isolate (Zika virus/H.sapiens-tc/PUR/2015/PRVABC59_v3c2) was provided by Brandy Russell (CDC, Fort Collins, CO, USA). On the day of the viral challenge, virus stocks were thawed, diluted to 1 mL with phosphate-buffered saline, and loaded into 1-mL syringes, which were kept on ice in a cooler until inoculation. Each dam was inoculated subcutaneously in the upper back, between the shoulder blades (cranial dorsum).

### Ultrasonographic analysis of fetal development during gestation

Ultrasounds were conducted every two weeks according to the procedures described previously [46] to monitor the heart rate and growth of the fetus. All dams were anesthetized with an intramuscular dose of ketamine (10 mg/kg) prior to all ultrasounds. Fetal measurements included fetal femur length (FL), biparietal diameter (BPD), head circumference (HC), and abdominal circumference (AC). All measurements were plotted against normative data collected by the California National Primate Research Center [55,56], containing mean measurements and standard deviations for specific gestational days of rhesus macaque pregnancy.

Z-scores represent the number of standard deviations a given fetal measurement deviates from the normative mean for that gestational age. For each metric (FL, BPD, HC, AC), we calculated Z-scores by comparing individual values to gestational age-matched reference data from the California National Primate Research Center [55,56]. To smooth variation in reference values across gestation, we fit quadratic regression models to the normative mean and standard deviation as functions of gestational age and used the resulting estimates to calculate predicted Z-scores. A one-sample *t*-test was used to determine whether the mean Z-score for each metric differed significantly from zero; in human fetal medicine, Z-scores less than 2 are generally not considered to be clinically meaningful [57,58]. To assess whether Z-scores changed over time, we used a linear mixed-effects model with animal-specific random effects and an autoregressive correlation structure, regressing Z-scores on gestational age. In this model, a significant slope indicated a systematic change in growth trajectory over gestation. Model assumptions were verified by examining residuals. All analyses were performed using SAS software (version 9.4; SAS Institute, Cary, NC), and two-sided *p*-values < 0.05 were considered statistically significant.

### Cesarean-section delivery and tissue collection

At ∼GD 155, all SPOV-inoculated pregnancies were surgically delivered via cesarean section, as previously described [41,46]. These were non-terminal survival procedures for the dams. We performed a comprehensive necropsy on the entire conceptus, including fetus, placental discs, fetal membranes, and umbilical cord. Fetuses were euthanized with an intravenous dose of at least 50 mg/kg sodium pentobarbital.

Tissue collection was performed under sterile conditions, as previously described [41,46]. Maternal biopsies were collected during surgery and included the spleen, liver, mesenteric lymph node, and uterine placental bed. Placental and fetal collections were performed immediately following cesarean section, as previously described [41,46]. Briefly, we collected a full-thickness center-cut of each placental disc and cotyledon. Individual cotyledons were then separated into decidua, chorionic plate, and parenchymal components. A full list of fetal, maternal, and maternal-fetal tissues collected can be found in **Supplemental Table 1**. Each tissue was visually examined and subsequently divided into sections for viral RNA quantification, histology, or long-term storage. When tissue sizes were limited, viral RNA quantification was prioritized.

### Processing of blood and urine samples

For larger blood draws, where we wanted to collect plasma and PBMC samples, we used the following protocol: EDTA-treated blood was layered over Ficoll in a 1:1 ratio using sterile transfer pipettes, centrifuged at 1860 relative centrifugal force (rcf; g-force) for 30 minutes, and brake set at 1. Sterile transfer pipettes were used to remove the plasma top layer into a sterile 15-mL conical tube. The “buffy coat” containing peripheral blood mononuclear cells (PBMCs) was then carefully transferred into a 15-mL conical tube with five milliliters of room-temperature R10 media. Both plasma and buffy coat tubes were spun at 670 rcf for 8 minutes. Plasma was stored in two 325-uL aliquots, with the remainder stored in larger aliquots. Media was removed from the buffy coat tubes, and the PBMC pellet was treated with 5 mL of ACK for 5 minutes, then quenched with 5 mL of R10. PBMCs were spun down at 670 rcf for 5 minutes, the media was removed, and the cells were resuspended in R10 before counting in the Beckman Coulter Counter. PBMCs were stored in 1 mL of CryoStor CS5 freezing media (STEMCELL Technologies, Cat #07930) in aliquots of 5e6–10e6 cells per tube. Samples were stored in Mr. Frosty containers (Thermo Fisher Scientific, Cat. No. 5100-0001) at -80°C overnight before moving to liquid nitrogen storage the next day.

For smaller blood draws, where we wanted to only collect plasma or serum, we used the following protocols: smaller, EDTA-treated blood samples were spun at 1400 rcf for 15 minutes. Plasma was transferred to a sterile tube with sterile transfer pipettes, then spun again at 670 rcf for 8 minutes. Plasma was stored in two 325-uL aliquots, with the remainder stored in larger aliquots. Blood in clot activator or serum-separating tubes was spun at 1400 rcf for 20 minutes, transferred to sterile tubes, and spun again at 670 rcf for 8 minutes. The serum was stored in 100 uL aliquots, with the remainder stored in larger aliquots. All plasma and serum samples were stored at -80°C.

Urine samples were collected via pan and were spun down at 500 rcf for 5 minutes to remove debris. 270 uL of urine, to be used for viral loads, was stored in 30 uL of DMSO and stored at 4°C before immediate testing. If the urine was not immediately tested, it was frozen in Mr. Frosty containers at -80°C. The remaining urine was stored in 1 mL aliquots at -80°C.

### Viral RNA isolation from fluids

Viral RNA was isolated using the Maxwell 48 RSC instrument (Promega, Madison WI) with the Maxwell Viral TNA kit (Promega, Madison, WI). Samples were vortexed and briefly spun to remove any sample from the tube lid. 330 uL of the provided lysis solution (from the provided lysis and proteinase K) was added to empty microcentrifuge tubes, then 300 uL of each sample was added. Each tube was vortexed and then briefly spun on a benchtop centrifuge before incubation on a 56°C heat block for 10 minutes. New gloves were donned, and the sample tray was prepared per the manufacturer’s instructions during the 10-minute incubation step.

Following incubation, all samples were briefly spun down and then run in the Maxwell 48 RSC instrument. After vRNA isolation, samples were transferred to new tubes and stored at -80°C if RT-PCR was performed later.

### Viral RNA isolation from tissues

Tissue samples were stored in RNAlater. RNA was recovered from tissue samples using a modification of the method described by Hansen et al. (2013) [69]. Briefly, up to 200 mg of tissue was disrupted in TRIzol (Lifetechnologies) with 2 x 5 mm stainless steel beads using the TissueLyser (Qiagen) for 3 minutes at 25 r/s twice. Following homogenization, samples in TRIzol were separated using Bromo-chloro-propane (Sigma). The aqueous phase was collected and glycogen was added as a carrier. The samples were washed in isopropanol and ethanol precipitated. RNA was fully re-suspended in 5 mM tris pH 8.0.

### Viral RNA quantification by RT-PCR

Quantitative reverse transcription-PCR (RT-PCR) was performed on a Roche LightCycler 480 (Roche Diagnostics, Indianapolis, IN) with ThermoFisher TaqMan Fast Virus 1-Step Master Mix for qPCR (Cat. No. 4444434) as previously described [52]. Briefly, samples were processed alongside water, where 10 uL of each RNA sample was added to water, master mix, 600 nM forward/reverse primers and random hexamers, and 100 nM probe. The plate containing the samples was sealed, spun for 10 seconds, and then run with the following cycling conditions: 50°C for 5 minutes (ramp rate: 4.4°C/second), 95°C for 20 seconds (ramp rate: 4.4°C/second), 50 cycles of 95°C for 15 seconds (ramp rate: 3.0°C/second) and 60°C for 1 minute (ramp rate: 2.2°C/second) before cooling at 40°C for 30 seconds (ramp rate: 2.2°C/second).

Primer and probe sequences for ZIKV RNA quantification:

Forward primer: 5’-CGYTGCCCAACACAAGG-3’ Reverse primer: 5’-CACYAAYGTTCTTTTGCABACAT-3’

Probe: 5’-6-carboxyfluorescein-AGCCTACCTTGAYAAGCARTCAGACACYCAA-BHQ1-3’

Primer and probe sequences for SPOV RNA quantification:

Forward primer: 5’-GGCATACAGGAGCCACATCAAAC-3’

Reverse primer: 5’-TGCGTGGGCTTCTCTGAA-3’

Probe: 5’-6-carboxyfluorescein-CATCACTGGAACAAYAAGGAGGCGCTGG-BHQ1-3’

The lower limit of detection (LLOD) of this assay in body fluids for ZIKV is 150 copies/ml and 81 copies/mg in tissues. The LLOD of this assay in body fluids for SPOV is 175 copies/ml and 108 copies/mg in tissues. LLOD is defined here as the lowest concentration of viral nucleic acid that can be reliably detected as true-positive with a 95% confidence interval.

### Quantification of plaque reduction neutralization by serum

Serum neutralization of SPOV was quantified using a plaque-reduction neutralization test (PRNT) on Vero cells, following the methods outlined by Jaeger *et al*. [52]. Serum samples were processed in clot activator or serum-separating tubes as described above and screened against the same SPOV stock used for viral inoculation. Neutralization curves were generated in GraphPad Prism 8 through nonlinear regression and evaluated using Prism’s ECanything equation to estimate EC50, which represents the serum dilution titer required to inhibit 50% of viral infectivity. Neutralization curves and EC50 values were generated for serum from each macaque at 0 and approximately 30 days post-inoculation.

### Histopathological analysis of placental, neural and ocular tissues

Tissue preparation and histopathological analysis were performed as previously described [46,59]. Briefly, placental tissues were fixed in 4% paraformaldehyde (PFA) overnight (∼24 hours) and then transferred to 70% ethanol until routine paraffin embedding. Neural and ocular tissues were fixed separately in 10% neutral buffered formalin (NBF), then routinely processed and paraffin embedded. Paraffin sections (5 μm) were cut and stained with hematoxylin and eosin (H&E) using standard protocols [46,59]. All histological assessments were conducted by veterinary pathologists under blinded conditions with respect to vRNA and fetal-growth findings.

### Data availability

Data, code, and figures are available for replication of these results on GitHub (https://github.com/RiesHunter/SPOV).

## Acknowledgments

We thank our colleagues at the Wisconsin National Primate Research Center (Animal Care Services, Veterinary Services, Pathology Services, and Scientific Protocol Implementation) for their invaluable care of the animals and their support throughout the study. We also thank members of the Friedrich and O’Connor laboratories for helpful discussions and technical assistance.

## Author contributions

**H.J.R.**—Conceptualization, Data Curation, Formal Analysis, Funding Acquisition, Investigation, Methodology, Project Administration, Software, Validation, Visualization, Writing – Original Draft Preparation, Writing – Review & Editing

**L.R.**—Data Curation, Investigation, Methodology, Validation, Writing – Review & Editing **M.C.C.**—Data Curation, Investigation, Methodology, Validation, Writing – Review & Editing

**C.M.C.**—Conceptualization, Data Curation, Funding Acquisition, Investigation, Methodology, Validation, Writing – Review & Editing

**C.D.**—Data Curation, Investigation, Methodology, Validation, Writing – Review & Editing

**A.R.**—Data Curation, Investigation, Methodology, Validation, Writing – Review & Editing

**G.V.S.**—Data Curation, Investigation, Methodology, Validation, Writing – Review & Editing

**A.M.W.**—Data Curation, Investigation, Methodology, Validation, Writing – Review & Editing

**J.C.E.**—Data Curation, Formal Analysis, Investigation, Methodology, Software, Validation, Writing – Review & Editing

**K.S.S.**—Data Curation, Formal Analysis, Investigation, Writing – Review & Editing

**L.B.C.T.**—Data Curation, Investigation, Writing – Review & Editing

**E.P.**—Investigation, Project Administration, Resources, Writing – Review & Editing

**M.S.**—Investigation, Project Administration, Resources, Writing – Review & Editing

**H.A.S.**—Data Curation, Formal Analysis, Investigation, Methodology, Resources, Validation, Writing – Review & Editing

**J.R.R.**—Investigation, Writing – Review & Editing

**L.E.R.**—Investigation, Writing – Review & Editing

**A.S.J.**—Conceptualization, Investigation, Methodology, Resources, Validation, Visualization, Writing – Review & Editing

**E.R.R.**—Investigation, Resources, Writing – Review & Editing

**E.L.M.**—Investigation, Resources, Writing – Review & Editing

**D.H.O.**—Investigation, Resources, Writing – Review & Editing

**C.M.N.**—Investigation, Resources, Writing – Review & Editing

**M.T.A.**—Conceptualization, Funding Acquisition, Project Administration, Resources, Supervision, Writing – Review & Editing

**T.C.F.**—Conceptualization, Funding Acquisition, Project Administration, Resources, Supervision, Validation, Writing – Original Draft Preparation, Writing – Review & Editing

## Additional information

Correspondence and requests for materials should be emailed to T.C.F.

## Supplemental Information

**S1 Fig.**
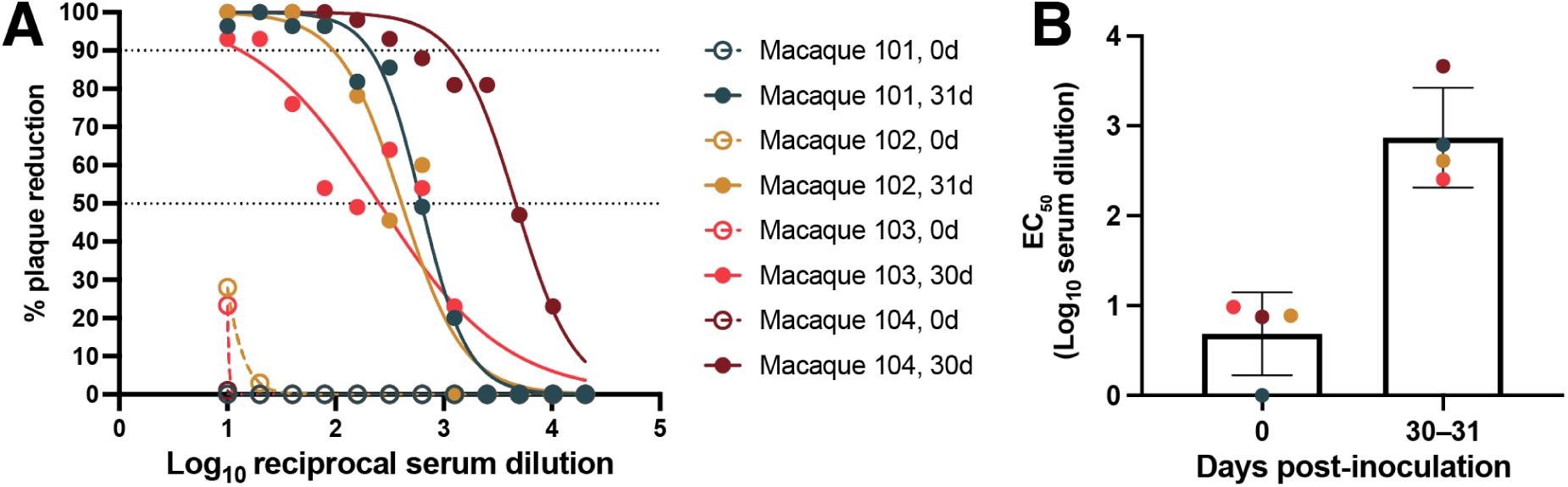
Neutralizing antibody responses to Spondweni virus at 0 and 30–31 days post-inoculation. **(A)** Plaque-reduction neutralization curves for each macaque at day 0 (pre-inoculation) and 30–31 days post-inoculation (dpi). The y-axis shows percent plaque reduction, and the x-axis shows log_10_ reciprocal serum dilution. The horizontal grey lines indicate 90% and 50% plaque reduction. **(B)** EC50 values (log_10_ serum dilution achieving 50% plaque reduction) for the same animals at 0 and 30–31 dpi. Points represent individual macaques (*n* = 4), and bars show group mean ± SD.

**S2 Fig.**
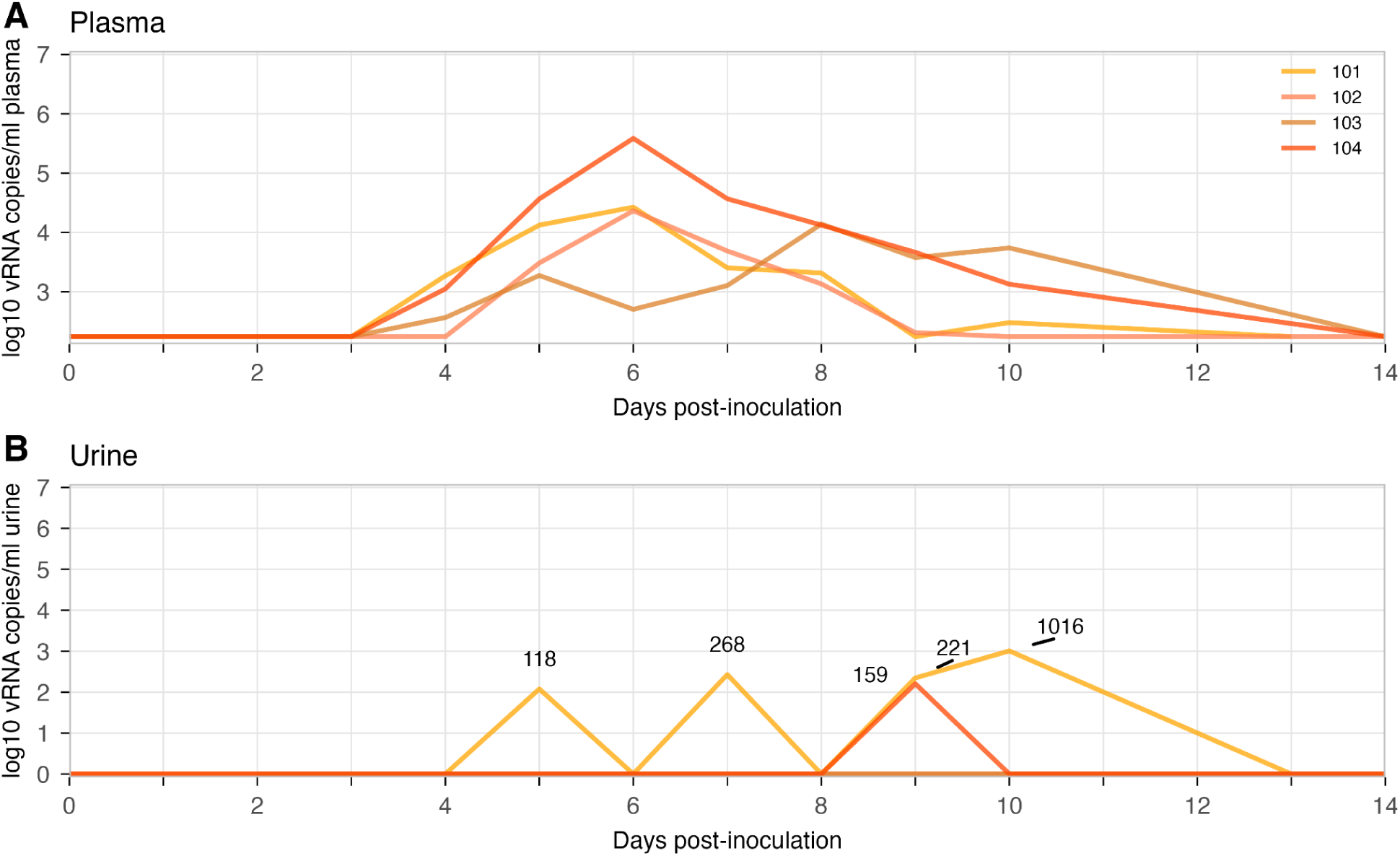
Acute-phase plasma viremia and sporadic urinary detection in pregnant macaques inoculated with Spondweni virus (SPOV). **(A)** Longitudinal plasma viral RNA (vRNA) loads (log10 copies/mL) in SPOV-inoculated pregnant rhesus macaques during the first 14 days post-inoculation (dpi). Lines trace individual animals. The x-axis is truncated at 14 days post-inoculation to emphasize the acute phase of infection and does not include the two plasma samples that tested positive at days 23 and 31 (see **Fig 2**). **(B)** Urine vRNA loads for the same animals over dpi 0–14. All virus-positive urine samples are labeled with their corresponding viral load values. Values of zero represent samples that tested negative in two technical replicates. The assay lower limit of detection for SPOV in urine is 175 copies/mL, and values below it (e.g., 118 and 159 copies/mL) are considered detectable but not reliably quantifiable. Abbreviations: SPOV, Spondweni virus; vRNA, viral RNA; dpi, days post inoculation; LLOD, lower limit of detection.

**S3 Fig.**
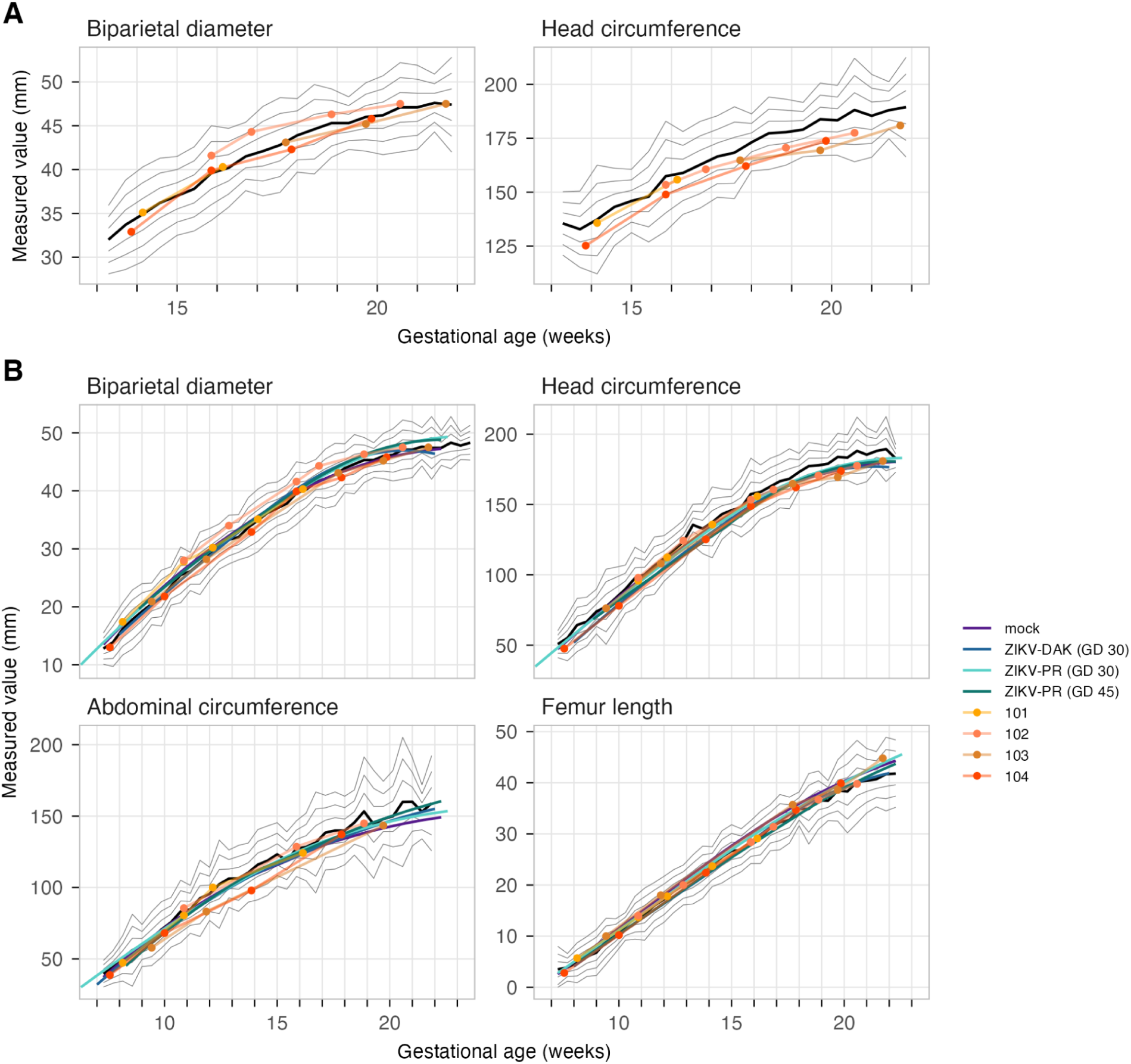
SPOV-exposed macaques’ fetal growth relative to normative curves and WNPRC cohorts. **(A)** Fetal growth trajectories for the SPOV-inoculated macaques’ biparietal diameter and head circumference from 13–22 weeks of gestation. Each point (*n* = 4) represents one macaque, with lines connecting individual macaques through time. Normative fetal measurements are shown as a black line (mean) with flanking grey lines representing ±1, ±2, and ±3 standard deviations at each gestational age. **(B)** Full dataset of fetal biparietal diameter, head circumference, abdominal circumference, and femur length. Fetal growth trajectory regressions are shown for mock-inoculated (purple), ZIKV-PR-inoculated (light blue), and ZIKV-DAK-inoculated (dark blue) macaques from prior studies [42].

**S4 Fig.**
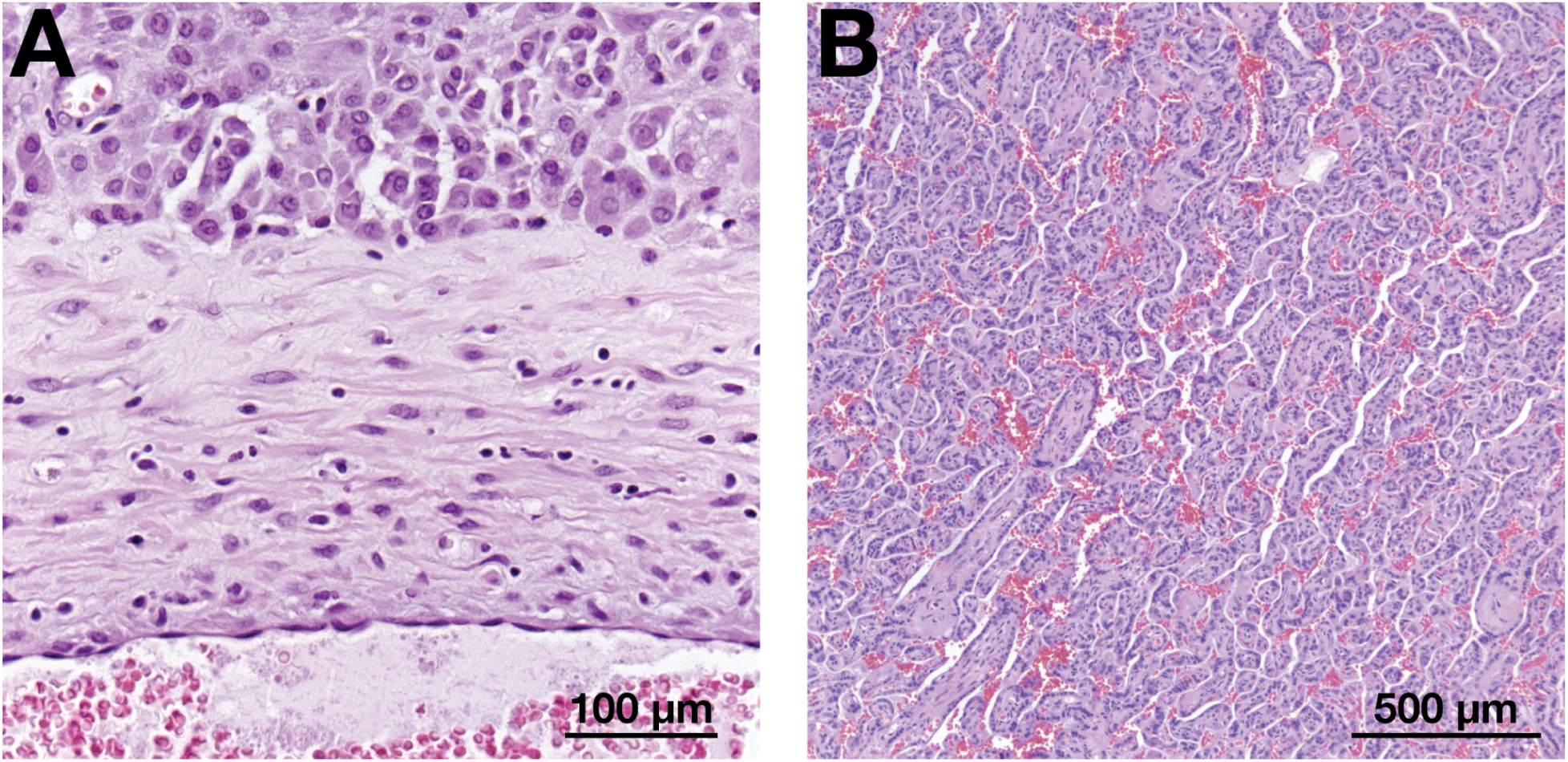
H&E-stained normal basal-plate artery and chorionic villous parenchyma from rhesus macaque placentas. **(A)** Maternal side of the placenta (basal plate) at 40× showing a normal spiral artery with a wide lumen and a thin vessel wall. No inflammation of the vessel wall (vasculitis) is present. Scale bar = 100 µm. **(B)** Chorionic villous parenchyma at 10× with preserved branching villi and open spaces between villi (intervillous spaces) containing red blood cells. No villitis, thrombi, perivillous fibrin, or infarction is evident. Scale bar = 500 µm. Panel A serves as a normal reference for Fig 5; Panel B serves as a normal reference for Fig 6. Abbreviations: H&E, hematoxylin and eosin.

**S5 Fig.**
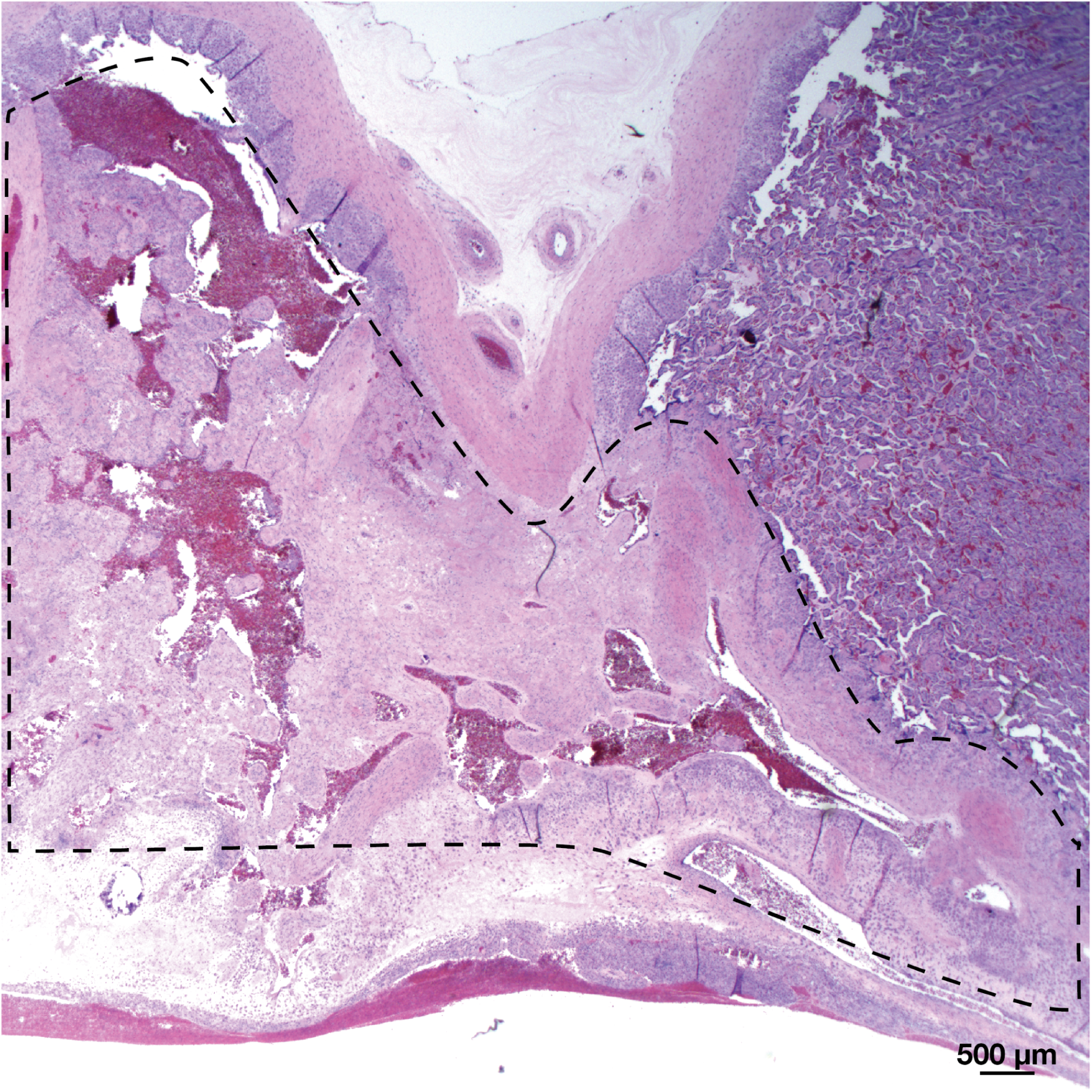
Representative image of transmural ischemia of the placenta cross-section from macaque 104. H&E-stained full-thickness placental section at delivery from macaque 104. The dashed outline marks ischemic placental parenchyma extending from the basal plate (bottom) to the chorionic plate (top), consistent with transmural placental ischemia. The chorionic plate has fallen into a “V shape” due to necrosis and loss of supporting villous tissue. Adjacent uninvolved chorionic villi are visible at right for comparison.

**Supplemental Table 1.** List of fetal, maternal, and maternal-fetal tissues collected during fetal necropsy near term.

| Fetal Tissues |  |
| --- | --- |
| CNS | Terminal CSF collection |
|  | Dura Mater |
|  | Cervical spinal cord |
|  | Thoracic spinal cord |
|  | Lumbar spinal cord |
|  | Cerebrum - right (10 sections) |
|  | Cerebrum - left |
|  | Cerebellum - 1 right-most median section |
|  | Cerebellum - left |
|  | Cerebellum - 3 right-most lateral section |
|  | Cerebellum - 2 right |
| Ocular | Aqueous humor - right aspirate and aliquot |
|  | Optic nerve - right |
|  | Eye - left |
|  | Sclera - right |
|  | Cornea - right |
|  | Retina - right |
| Cardiopulmonary | Pericardium |
|  | Heart full thickness section |
|  | Aorta - thoracic |
|  | Lung |
| Reproductive | Sem vesicle & prostate/uterus |
|  | Testis/Ovary |
| Musculoskeletal | Adipose tissue - omentum |
|  | Epidermis/dermis of abdomen |
|  | Muscle - quadriceps |
| Immune | Bone marrow - if possible R femur only |
|  | Tonsil oropharyngeal LN |
|  | Spleen |
|  | Thymus |
|  | Submandibular LN |
|  | Tracheobronchial LN |
|  | Mesenteric LN |
|  | Axillary LN |
|  | Inguinal LN |
| Gastrointestinal | Esophagus |
|  | Stomach |
|  | Duodenum |
|  | Jejunum |
|  | Ileum |
|  | Cecum |
|  | Colon |
|  | Liver |
|  | Tongue |
| Urinary | Urinary bladder |
|  | Kidney - right and left |
|  | Urine - aspirate |
| Endocrine | Thyroid |
|  | Adrenal Gland |
|  | Pancreas |

| Maternal tissues |
| --- |
| Spleen |
| Liver |
| Mesenteric LN |
| Uterus - Placental bed (point of attachment) |

| Maternal-fetal interface |
| --- |
| Amniotic fluid |
| Umbilical cord blood |
| Chorionic plate |
| Fetal membranes (taken distant from placenta) |
| Placenta disc 2 full thickness |
| Placental disc 1 periphery - no decidua |
| Placental disc 2 periphery - no decidua |
| Decidua - removed from placenta (Basalis) |
| Inter-placental collateral vessels |
| Umbilical cord |

**Supplemental Table 2.**
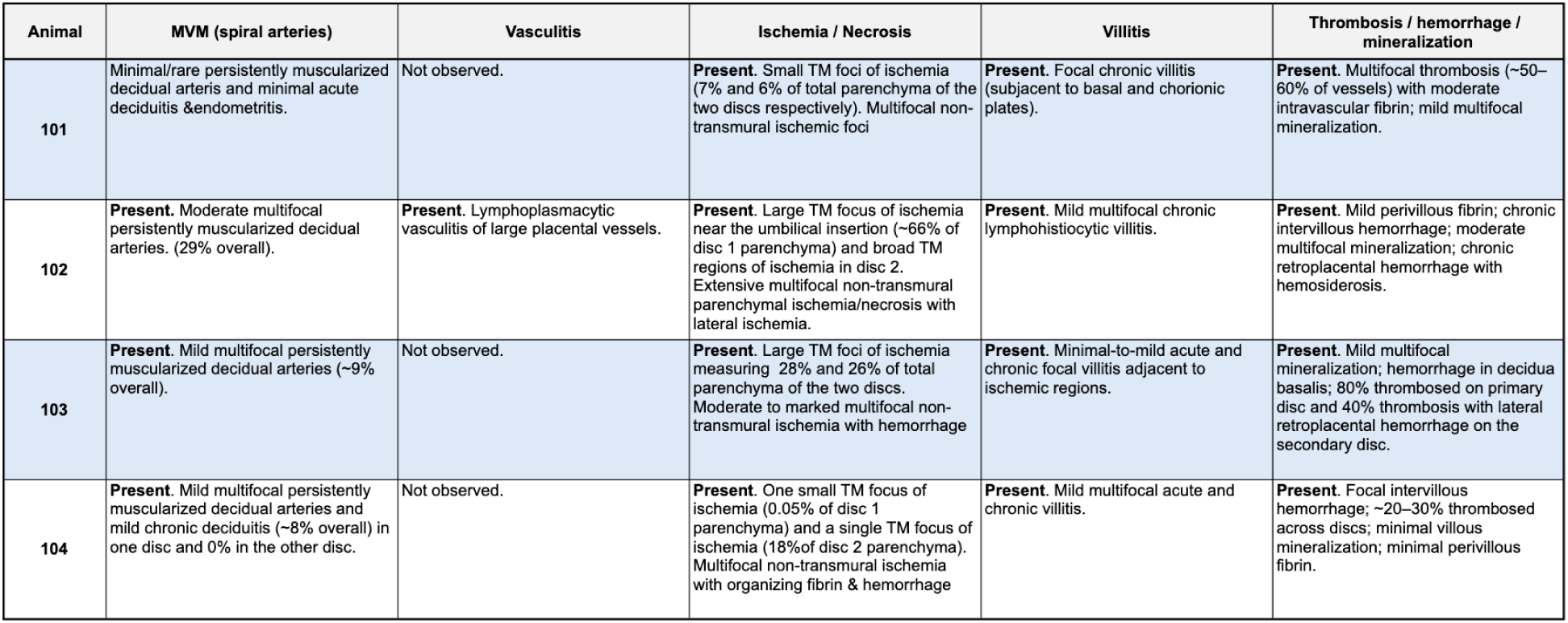
Placental histopathology by animal. H&E-stained placental sections collected at delivery were reviewed for two discs per pregnancy (center cut plus cotyledon samples). Cells list the dominant findings across both discs for each animal. Abbreviations: MVM = maternal vascular malperfusion and TM = transmural.

| Animal | MVM (spiral arteries) | Vasculitis | Ischemia / Necrosis | Villitis | Thrombosis / hemorrhage / mineralization |
| --- | --- | --- | --- | --- | --- |
| 101 | Minimal/rare persistently muscularized decidual arteries and minimal acute deciduitis & endometritis. | Not observed. | **Present.** Small TM foci of ischemia (7% and 6% of total parenchyma of the two discs respectively). Multifocal non-transmural ischemic foci | **Present.** Focal chronic villitis (subjacent to basal and chorionic plates). | **Present.** Multifocal thrombosis (~50–60% of vessels) with moderate intravascular fibrin; mild multifocal mineralization. |
| 102 | **Present.** Moderate multifocal persistently muscularized decidual arteries. (29% overall). | **Present.** Lymphoplasmacytic vasculitis of large placental vessels. | **Present.** Large TM focus of ischemia near the umbilical insertion (~66% of disc 1 parenchyma) and broad TM regions of ischemia in disc 2. Extensive multifocal non-transmural parenchymal ischemia/necrosis with lateral ischemia. | **Present.** Mild multifocal chronic lymphohistiocytic villitis. | **Present.** Mild perivillous fibrin; chronic intervillous hemorrhage; moderate multifocal mineralization; chronic retroplacental hemorrhage with hemosiderosis. |
| 103 | **Present.** Mild multifocal persistently muscularized decidual arteries (~9% overall). | Not observed. | **Present.** Large TM foci of ischemia measuring 28% and 26% of total parenchyma of the two discs. Moderate to marked multifocal non-transmural ischemia with hemorrhage | **Present.** Minimal-to-mild acute and chronic focal villitis adjacent to ischemic regions. | **Present.** Mild multifocal mineralization; hemorrhage in decidua basalis; 80% thrombosed on primary disc and 40% thrombosis with lateral retroplacental hemorrhage on the secondary disc. |
| 104 | **Present.** Mild multifocal persistently muscularized decidual arteries and mild chronic deciduitis (~8% overall) in one disc and 0% in the other disc. | Not observed. | **Present.** One small TM focus of ischemia (0.05% of disc 1 parenchyma) and a single TM focus of ischemia (18% of disc 2 parenchyma). Multifocal non-transmural ischemia with organizing fibrin & hemorrhage | **Present.** Mild multifocal acute and chronic villitis. | **Present.** Focal intervillous hemorrhage; ~20–30% thrombosed across discs; minimal villous mineralization; minimal perivillous fibrin. |

## References

1. Pierson TC, Diamond MS. The continued threat of emerging flaviviruses. Nat Microbiol. 2020;5: 796–812. doi:10.1038/s41564-020-0714-0

2. Bhatt S, Gething PW, Brady OJ, Messina JP, Farlow AW, Moyes CL, et al. The global distribution and burden of dengue. Nature. 2013;496: 504–507. doi:10.1038/nature12060

3. Simmonds P, Becher P, Bukh J, Gould EA, Meyers G, Monath T, et al. ICTV Virus Taxonomy Profile: Flaviviridae. J Gen Virol. 2017;98: 2–3. doi:10.1099/jgv.0.000672

4. Brady OJ, Gething PW, Bhatt S, Messina JP, Brownstein JS, Hoen AG, et al. Refining the Global Spatial Limits of Dengue Virus Transmission by Evidence-Based Consensus. PLoS Negl Trop Dis. 2012;6: e1760. doi:10.1371/journal.pntd.0001760

5. Ryan SJ, Carlson CJ, Mordecai EA, Johnson LR. Global expansion and redistribution of Aedes-borne virus transmission risk with climate change. PLoS Negl Trop Dis. 2019;13: e0007213. doi:10.1371/journal.pntd.0007213

6. Vaughn DW, Green S, Kalayanarooj S, Innis BL, Nimmannitya S, Suntayakorn S, et al. Dengue in the early febrile phase: viremia and antibody responses. J Infect Dis. 1997;176: 322–330. doi:10.1086/514048

7. Yacoub S, Wertheim H, Simmons CP, Screaton G, Wills B. Microvascular and endothelial function for risk prediction in dengue: an observational study. Lancet Lond Engl. 2015;385 Suppl 1: S102. doi:10.1016/S0140-6736(15)60417-2

8. CDC. Clinical Features of Dengue. In: Dengue [Internet]. 15 May 2025 [cited 3 Sept 2025]. Available: https://www.cdc.gov/dengue/hcp/clinical-signs/index.html

9. Quaresma JAS, Pagliari C, Medeiros DBA, Duarte MIS, Vasconcelos PFC. Immunity and immune response, pathology and pathologic changes: progress and challenges in the immunopathology of yellow fever. Rev Med Virol. 2013;23: 305–318. doi:10.1002/rmv.1752

10. CDC. Yellow Fever. In: Yellow Book [Internet]. 23 June 2025 [cited 3 Sept 2025]. Available: https://www.cdc.gov/yellow-book/hcp/travel-associated-infections-diseases/yellow-fever.html

11. PAHO. 1 December 2015: Neurological syndrome, congenital malformations, and Zika virus infection. Implications for public health in the Americas – Epidemiological Alert - PAHO/WHO | Pan American Health Organization. 1 Dec 2015 [cited 18 June 2025]. Available: https://www.paho.org/en/documents/1-december-2015-neurological-syndrome-congenital-malformations-and-zika-virus-infection

12. Hoen B, Schaub B, Funk AL, Ardillon V, Boullard M, Cabié A, et al. Pregnancy Outcomes after ZIKV Infection in French Territories in the Americas. N Engl J Med. 2018;378: 985–994. doi:10.1056/NEJMoa1709481

13. Shapiro-Mendoza CK. Pregnancy Outcomes After Maternal Zika Virus Infection During Pregnancy — U.S. Territories, January 1, 2016–April 25, 2017. MMWR Morb Mortal Wkly Rep. 2017;66. doi:10.15585/mmwr.mm6623e1

14. Reynolds MR. Vital Signs: Update on Zika Virus–Associated Birth Defects and Evaluation of All U.S. Infants with Congenital Zika Virus Exposure — U.S. Zika Pregnancy Registry, 2016. MMWR Morb Mortal Wkly Rep. 2017;66. doi:10.15585/mmwr.mm6613e1

15. Dick GWA, Kitchen SF, Haddow AJ. Zika Virus (I). Isolations and serological specificity. Trans R Soc Trop Med Hyg. 1952;46: 509–520. doi:10.1016/0035-9203(52)90042-4

16. Musso D, Gubler DJ. Zika Virus. Clin Microbiol Rev. 2016;29: 487–524. doi:10.1128/CMR.00072-15

17. Duffy MR, Chen T-H, Hancock WT, Powers AM, Kool JL, Lanciotti RS, et al. Zika virus outbreak on Yap Island, Federated States of Micronesia. N Engl J Med. 2009;360: 2536–2543. doi:10.1056/NEJMoa0805715

18. Cao-Lormeau V-M, Roche C, Teissier A, Robin E, Berry A-L, Mallet H-P, et al. Zika Virus, French Polynesia, South Pacific, 2013. Emerg Infect Dis. 2014;20: 1085–1086. doi:10.3201/eid2006.140138

19. Kleber de Oliveira W, Cortez-Escalante J, De Oliveira WTGH, do Carmo GMI, Henriques CMP, Coelho GE, et al. Increase in Reported Prevalence of Microcephaly in Infants Born to Women Living in Areas with Confirmed Zika Virus Transmission During the First Trimester of Pregnancy - Brazil, 2015. MMWR Morb Mortal Wkly Rep. 2016;65: 242–247. doi:10.15585/mmwr.mm6509e2

20. Teixeira MG, Costa M da CN, de Oliveira WK, Nunes ML, Rodrigues LC. The Epidemic of Zika Virus-Related Microcephaly in Brazil: Detection, Control, Etiology, and Future Scenarios. Am J Public Health. 2016;106: 601–605. doi:10.2105/AJPH.2016.303113

21. Moore CA, Staples JE, Dobyns WB, Pessoa A, Ventura CV, Fonseca EB da, et al. Characterizing the Pattern of Anomalies in Congenital Zika Syndrome for Pediatric Clinicians. JAMA Pediatr. 2017;171: 288–295. doi:10.1001/jamapediatrics.2016.3982

22. Massetti T, Herrero D, Alencar J, Silva T, Moriyama C, Gehrke F, et al. Clinical characteristics of children with congenital Zika syndrome: a case series. Arq Neuropsiquiatr. 2020;78: 403–411. doi:10.1590/0004-282X20200020

23. Zika Virus Individual Participant Data Consortium. Adverse fetal and perinatal outcomes associated with Zika virus infection during pregnancy: an individual participant data meta-analysis. EClinicalMedicine. 2025;83: 103231. doi:10.1016/j.eclinm.2025.103231

24. Haddow AD, Nasar F, Guzman H, Ponlawat A, Jarman RG, Tesh RB, et al. Genetic Characterization of Spondweni and Zika Viruses and Susceptibility of Geographically Distinct Strains of Aedes aegypti, Aedes albopictus and Culex quinquefasciatus (Diptera: Culicidae) to Spondweni Virus. PLoS Negl Trop Dis. 2016;10: e0005083. doi:10.1371/journal.pntd.0005083

25. de Groot R, Cowley J, Enjuanes L, Faaberg K, Perlman S, Rottier P, et al. ICTV 9th Report (2011) | ICTV. 2012 [cited 4 June 2025]. Available: https://ictv.global/report_9th

26. Theiler M, Downs WG. The arthropod-borne viruses of vertebrates: an account of the Rockefeller Foundation Virus Program, 1951-1970. New Haven: Yale University Press, 1973; 1973.

27. Haddow AD, Woodall JP. Distinguishing between Zika and Spondweni viruses. Bull World Health Organ. 2016;94: 711–711A. doi:10.2471/BLT.16.181503

28. Rathore APS, St John AL. Cross-Reactive Immunity Among Flaviviruses. Front Immunol. 2020;11: 334. doi:10.3389/fimmu.2020.00334

29. Macnamara FN. Zika virus: a report on three cases of human infection during an epidemic of jaundice in Nigeria. Trans R Soc Trop Med Hyg. 1954;48: 139–145. doi:10.1016/0035-9203(54)90006-1

30. Bearcroft WG. Zika virus infection experimentally induced in a human volunteer. Trans R Soc Trop Med Hyg. 1956;50: 442–448.

31. Mcintosh BM, Kokernot RH, Paterson HE, De Meillon B. Isolation of Spondweni virus from four species of culicine mosquitoes and a report of two laboratory infections with the virus. South Afr Med J Suid-Afr Tydskr Vir Geneeskd. 1961;35: 647–650.

32. Draper CC. INFECTION WITH THE CHUKU STRAIN OF SPONDWENI VIRUS. West Afr Med J. 1965;14: 16–19.

33. Wolfe MS, Calisher CH, McGuire K. Spondweni virus infection in a foreign resident of Upper Volta. Lancet Lond Engl. 1982;2: 1306–1308. doi:10.1016/s0140-6736(82)91511-2

34. Haddow AJ, Williams MC, Woodall JP, Simpson DIH, Goma LKH. Twelve Isolations of Zika Virus from Aedes (Stegomyla) africanus (Theobald) taken in and above a Uganda Forest.

35. Brottes H, Rickenbach A, Brès P, Salaün JJ, Ferrara L. [Arboviruses in the Cameroon. Isolation from mosquitoes]. Bull World Health Organ. 1966;35: 811–825.

36. White SK, Lednicky JA, Okech BA, Morris JG, Dunford JC. Spondweni Virus in Field-Caught Culex quinquefasciatus Mosquitoes, Haiti, 2016. Emerg Infect Dis. 2018;24: 1765–1767. doi:10.3201/eid2409.171957

37. Jaeger AS, Weiler AM, Moriarty RV, Rybarczyk S, O’Connor SL, O’Connor DH, et al. Spondweni virus causes fetal harm in Ifnar1−/− mice and is transmitted by Aedes aegypti mosquitoes. Virology. 2020;547: 35–46. doi:10.1016/j.virol.2020.05.005

38. Carter AM. Animal models of human placentation--a review. Placenta. 2007;28 Suppl A: S41–47. doi:10.1016/j.placenta.2006.11.002

39. Bohm EK, Vangorder-Braid JT, Jaeger AS, Moriarty RV, Baczenas JJ, Bennett NC, et al. Zika Virus Infection of Pregnant Ifnar1-/- Mice Triggers Strain-Specific Differences in Fetal Outcomes. J Virol. 2021;95: e0081821. doi:10.1128/JVI.00818-21

40. Raasch LE, Yamamoto K, Newman CM, Rosinski JR, Shepherd PM, Razo E, et al. Fetal loss in pregnant rhesus macaques infected with high-dose African-lineage Zika virus. PLoS Negl Trop Dis. 2022;16: e0010623. doi:10.1371/journal.pntd.0010623

41. Rosinski JR, Raasch LE, Barros Tiburcio P, Breitbach ME, Shepherd PM, Yamamoto K, et al. Frequent first-trimester pregnancy loss in rhesus macaques infected with African-lineage Zika virus. PLoS Pathog. 2023;19: e1011282. doi:10.1371/journal.ppat.1011282

42. Crooks CM, Weiler AM, Rybarczyk SL, Bliss M, Jaeger AS, Murphy ME, et al. African-Lineage Zika Virus Replication Dynamics and Maternal-Fetal Interface Infection in Pregnant Rhesus Macaques. J Virol. 2021;95: e0222020. doi:10.1128/JVI.02220-20

43. Crooks CM, Weiler AM, Rybarczyk SL, Bliss MI, Jaeger AS, Murphy ME, et al. Previous exposure to dengue virus is associated with increased Zika virus burden at the maternal-fetal interface in rhesus macaques. PLoS Negl Trop Dis. 2021;15: e0009641. doi:10.1371/journal.pntd.0009641

44. Aliota MT, Dudley DM, Newman CM, Mohr EL, Gellerup DD, Breitbach ME, et al. Heterologous Protection against Asian Zika Virus Challenge in Rhesus Macaques. PLoS Negl Trop Dis. 2016;10: e0005168. doi:10.1371/journal.pntd.0005168

45. Dudley DM, Aliota MT, Mohr EL, Weiler AM, Lehrer-Brey G, Weisgrau KL, et al. A rhesus macaque model of Asian-lineage Zika virus infection. Nat Commun. 2016;7: 12204. doi:10.1038/ncomms12204

46. Nguyen SM, Antony KM, Dudley DM, Kohn S, Simmons HA, Wolfe B, et al. Highly efficient maternal-fetal Zika virus transmission in pregnant rhesus macaques. PLoS Pathog. 2017;13: e1006378. doi:10.1371/journal.ppat.1006378

47. The Macaque Placenta—A Mini-Review - Eveline P. C. T. de Rijk, Eric Van Esch, 2008. [cited 18 June 2025]. Available: https://journals.sagepub.com/doi/10.1177/0192623308326095

48. Newman C, Friedrich TC, O’Connor DH. Macaque monkeys in Zika virus research: 1947-present. Curr Opin Virol. 2017;25: 34–40. doi:10.1016/j.coviro.2017.06.011

49. Li A, Coffey LL, Mohr EL, Raper J, Chahroudi A, Ausderau KK, et al. Role of non-human primate models in accelerating research and developing countermeasures against Zika virus infection. Lancet Microbe. 2025;6: 101030. doi:10.1016/j.lanmic.2024.101030

50. Koenig MR, Razo E, Mitzey A, Newman CM, Dudley DM, Breitbach ME, et al. Quantitative definition of neurobehavior, vision, hearing and brain volumes in macaques congenitally exposed to Zika virus. PloS One. 2020;15: e0235877. doi:10.1371/journal.pone.0235877

51. Ausderau KK, Boerigter B, Razo ER, Gutkes J, Krabbe NP, Mitzey AM, et al. Prenatal Zika virus exposure disrupts social-emotional development and cortical visual function in infant macaques. Nat Commun. 2026;17: 1803. doi:10.1038/s41467-026-68517-x

52. Jaeger AS, Crooks CM, Weiler AM, Bliss MI, Rybarczyk S, Richardson A, et al. Primary infection with Zika virus provides one-way heterologous protection against Spondweni virus infection in rhesus macaques. Sci Adv. 2023;9: eadg3444. doi:10.1126/sciadv.adg3444

53. Krabbe NP, Razo E, Abraham HJ, Spanton RV, Shi Y, Bhattacharya S, et al. Control of maternal Zika virus infection during pregnancy is associated with lower antibody titers in a macaque model. Front Immunol. 2023;14: 1267638. doi:10.3389/fimmu.2023.1267638

54. Styer LM, Kent KA, Albright RG, Bennett CJ, Kramer LD, Bernard KA. Mosquitoes inoculate high doses of West Nile virus as they probe and feed on live hosts. PLoS Pathog. 2007;3: 1262–1270. doi:10.1371/journal.ppat.0030132

55. Tarantal AF, Hendrickx AG. Prenatal growth in the cynomolgus and rhesus macaque (Macaca fascicularis and Macaca mulatta): A comparison by ultrasonography. Am J Primatol. 1988;15: 309–323. doi:10.1002/ajp.1350150405

56. Tarantal F. Ultrasound Imaging in Rhesus (Macaca Mulatta) and Long-Tailed (Macaca fascicularis) Macaques. Reproductive and Research Applications. The laboratory primate. Amsterdam, The Netherlands: In Wolfe-Coote S (ed), Elsevier Ltd; 2005. pp. 317–352. doi:10.1016/B978-012080261-6/50020-9

57. Salomon LJ, Alfirevic Z, Da Silva Costa F, Deter RL, Figueras F, Ghi T, et al. ISUOG Practice Guidelines: ultrasound assessment of fetal biometry and growth. Ultrasound Obstet Gynecol Off J Int Soc Ultrasound Obstet Gynecol. 2019;53: 715–723. doi:10.1002/uog.20272

58. Society for Maternal-Fetal Medicine (SMFM) Publications Committee. Ultrasound screening for fetal microcephaly following Zika virus exposure. Am J Obstet Gynecol. 2016;214: B2–4. doi:10.1016/j.ajog.2016.02.043

59. Mohr EL, Block LN, Newman CM, Stewart LM, Koenig M, Semler M, et al. Ocular and uteroplacental pathology in a macaque pregnancy with congenital Zika virus infection. PloS One. 2018;13: e0190617. doi:10.1371/journal.pone.0190617

60. Hirsch AJ, Roberts VHJ, Grigsby PL, Haese N, Schabel MC, Wang X, et al. Zika virus infection in pregnant rhesus macaques causes placental dysfunction and immunopathology. Nat Commun. 2018;9: 263. doi:10.1038/s41467-017-02499-9

61. Dudley DM, Koenig MR, Stewart LM, Semler MR, Newman CM, Shepherd PM, et al. Human immune globulin treatment controls Zika viremia in pregnant rhesus macaques. PloS One. 2022;17: e0266664. doi:10.1371/journal.pone.0266664

62. Koenig MR, Mitzey AM, Morgan TK, Zeng X, Simmons HA, Mejia A, et al. Infection of the maternal-fetal interface and vertical transmission following low-dose inoculation of pregnant rhesus macaques (Macaca mulatta) with an African-lineage Zika virus. PloS One. 2023;18: e0284964. doi:10.1371/journal.pone.0284964

63. Koenig MR, Mitzey AM, Zeng X, Reyes L, Simmons HA, Morgan TK, et al. Vertical transmission of African-lineage Zika virus through the fetal membranes in a rhesus macaque (Macaca mulatta) model. PLoS Pathog. 2023;19: e1011274. doi:10.1371/journal.ppat.1011274

64. Szaba FM, Tighe M, Kummer LW, Lanzer KG, Ward JM, Lanthier P, et al. Zika virus infection in immunocompetent pregnant mice causes fetal damage and placental pathology in the absence of fetal infection. PLoS Pathog. 2018;14: e1006994. doi:10.1371/journal.ppat.1006994

65. Nielsen-Saines K, Brasil P, Kerin T, Vasconcelos Z, Gabaglia CR, Damasceno L, et al. Delayed childhood neurodevelopment and neurosensory alterations in the second year of life in a prospective cohort of ZIKV-exposed children. Nat Med. 2019;25: 1213–1217. doi:10.1038/s41591-019-0496-1

66. Mulkey SB, Arroyave-Wessel M, Peyton C, Bulas DI, Fourzali Y, Jiang J, et al. Neurodevelopmental Abnormalities in Children With In Utero Zika Virus Exposure Without Congenital Zika Syndrome. JAMA Pediatr. 2020;174: 269–276. doi:10.1001/jamapediatrics.2019.5204

67. Dudley DM, Newman CM, Lalli J, Stewart LM, Koenig MR, Weiler AM, et al. Infection via mosquito bite alters Zika virus tissue tropism and replication kinetics in rhesus macaques. Nat Commun. 2017;8: 2096. doi:10.1038/s41467-017-02222-8

68. The Weatherall report on the use of non-human primates in research | Royal Society. [cited 17 June 2025]. Available: https://royalsociety.org/news-resources/publications/2006/weatherall-report/

69. Hansen SG, Piatak M, Ventura AB, Hughes CM, Gilbride RM, Ford JC, et al. Immune clearance of highly pathogenic SIV infection. Nature. 2013;502: 100–104. doi:10.1038/nature12519

